# Cell atlases of schistosome human infective stages reveal larval tissue fates and innovations

**DOI:** 10.64898/2026.11.13.766466

**Authors:** Teresa Attenborough, Sarah O. Kerr, Yuxuan Xu, Frances Blow, Magda Lotkowska, Jennie Graham, Simon Kershenbaum, Gabriel Rinaldi, Matthew Berriman, Kate Rawlinson

## Abstract

Schistosomes infect over 250 million people, yet the cellular basis of their human-infective cercaria larva, which drives transmission and infection, remains poorly understood. We present single-cell atlases of *Schistosoma mansoni* cercariae, considered transcriptionally quiescent, and schistosomula, the first intramammalian stage. Across 22 spatially validated cell types, we identify populations persisting into the schistosomulum and populations lost. The tail harbours cercaria-specific innovations, including a specialised tegument. Gene-age analyses support the cercaria as the most recent addition to the life cycle and tail cells as younger than body cells. Persisting cell types undergo clear transcriptional transitions, providing insights into the dynamics of cercarial transformation into schistosomula. These findings link the recent origin of cercarial cell types to the evolution of active dispersal and transmission in digenean parasites.

## Main text

Schistosomes, which infect more than 250 million people *(1)*, are blood flukes with a complex life cycle involving a molluscan intermediate host, a mammalian definitive host and two free-swimming larval stages in between (fig. S1A). No vaccine exists, and praziquantel, the only available drug, is active only against adult worms. With frequent reinfection and growing concern over drug resistance, new interventions, particularly those targeting transmission, are urgently needed.

Transmission to humans depends on the cercaria, the human-infective larva. Cercariae emerge from the snail host and swim with their forked tail. They do not feed and their energy reserves last up to 53 hours, so within this time they must locate and penetrate human skin, but infectivity declines rapidly 8 hours post-emergence *(2, 3)*. On invasion the tail is shed, and the body transforms into the schistosomulum, which enters the vasculature and ultimately matures into an adult worm *(4, 5)* (fig. S1A). The cercaria is an evolutionary novelty. Tailed, non-feeding and gland-rich, it is found nowhere outside Digenea and is thought to be the most recent addition to the life cycle *(6)*. Its tail-driven, active transmission likely replaced the passive infection of vertebrates through ingestion of infected molluscs *(7)*.

To establish infection while transitioning to a parasitic form and adapting to a completely different environment, the cercaria replaces its gene expression programme with that of the schistosomulum. Cercariae are described as transcriptionally silent, producing the transcripts they need before leaving the snail *(8, 9)*. This view largely rests on sparse nascent-RNA labelling of whole cercariae after 18h incubation, and bivalent chromatin marks at 121 genes, which may poise them for transcriptional reactivation after transformation *(10)*. Whether quiescence extends to all cell types, and how cercarial transcripts are cleared after transformation, are unknown.

Single-cell transcriptomics has recently resolved several schistosome life stages at cellular resolution *(11–14)*. However, the cercaria and its transition into the schistosomulum remain poorly characterized.

We do not know the full diversity of cercarial cell types. Which of them continue into the schistosomulum, and which of them are unique to the infective larva? Are unique cell types limited to the tail that is shed during infection? These gaps limit our understanding of cell types and molecular mechanisms that drive transmission and host invasion. We generated a single-cell atlas of *Schistosoma mansoni* cercariae, captured within four hours of emergence from the snail, alongside a new 2-day schistosomula atlas, more deeply sampled than the published version *(12)*, generated with matched methods to allow direct comparison across the transition (fig. S1B). These allow us to define the cercarial cell types and their spatial organisation, use cell-type-restricted genes to infer the evolutionary origins of cercarial cell types, and characterise the cellular transcriptional programmes involved in the transition to the intramammalian environment.

The cercarial atlas comprises 24,708 cells, resolved into 24 transcriptionally distinct, mixed-sex populations covering all major tissues except the acetabular glands, which were too large for droplet capture (Fig. 1A and fig. S1C; table S1). We annotated clusters using published *S. mansoni* markers (data S2), SAMap comparisons with other developmental stages and other flatworms (fig. S2; data S2) *(15–17)*, and whole-mount *in situ* hybridization (ISH) of top marker genes (Fig. 1, B to F, and fig. S3; table S2). ISH spatially validated 22 clusters (figs. S4 to S10); 11 cell types were found in the body only, five in both body and tail, and six in the tail only (fig. S11; table S3). Each cell type is described in detail in the supplementary text; expression data and interactive three-dimensional reconstructions of the ISH images can be explored in an online atlas (schistoviewer.org).

**Fig 1.**
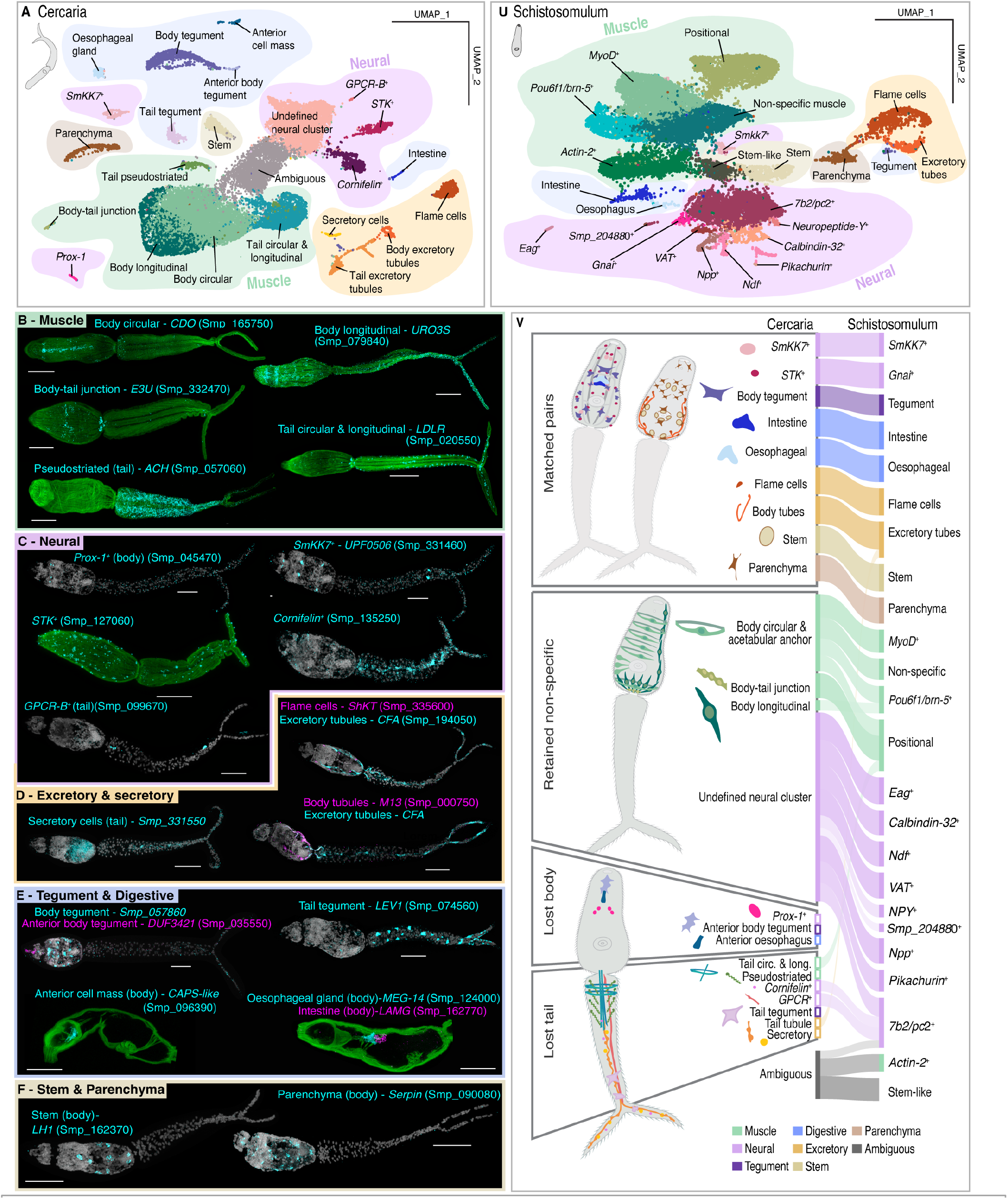
Single-cell atlas of *Schistosoma mansoni* cercaria and schistosomulum. **(A)** UMAP plot of the 24 scRNA-seq clusters of the cercaria. **(B to F)** *In situ* hybridization of marker genes of 22 cercarial cell type clusters. Phalloidin-labelled actin (green), DAPI (gray). Scale bars, 50 µm. **(G)** UMAP plot of the 24 scRNA-seq clusters of the schistosomulum. **(H)** SAMap alignment of cercarial cell types to schistosomulum clusters revealed nine clusters that pair 1:1, four that align with multiple clusters, three in the cercarial body that do not match schistosomulum clusters and seven that are enriched *(Cornifelin*+ neurons) or specific to the tail and are lost with the tail. **2. Cell type architecture of the cercarial tail. (A)** Nuclear segmentation: 832 nuclei in the body and 285 in the tail. **(B)** 3D rendering of tail muscle marker expression. **(C)** Segmented nuclei of tail neuronal cell types. **(Di)** Dextran labelling shows cilia of tail, including ciliary loops (▾). **(Dii to Div)** Scanning electron micrographs (SEM) show ciliary morphologies: clubbed (Dii and Diii) and looped clubbed (Div). **(E)** 3D rendering of excretory cell type markers expression. **(F)** Segmented secretory cell marker expression, also in the body acetabular glands (arrow). **(Gi)** 3D rendering of tail tegument marker, *LEV1*, expression, **(Gii)** and actual expression in large domains along the tail. **(Giii)** Cytoplasmic protrusions of *LEV1*^+^ cells (arrows) extend between muscles (phalloidin; green). **(Hi)** SEM of a cercaria, with body (Hii) and tail (Hiii) tegument spines magnified. **(I)** Tail tegument spines are larger and more widely spaced than body spines; *n* = 30 spines per region; Welch two-sample *t* tests, *P* < 0.005. Asterisks, cell types only in the tail. All scale bars, 50 µm except; Dii to Div, 1 µm; Hii and Hiii, 2 µm.

**Fig 2.**
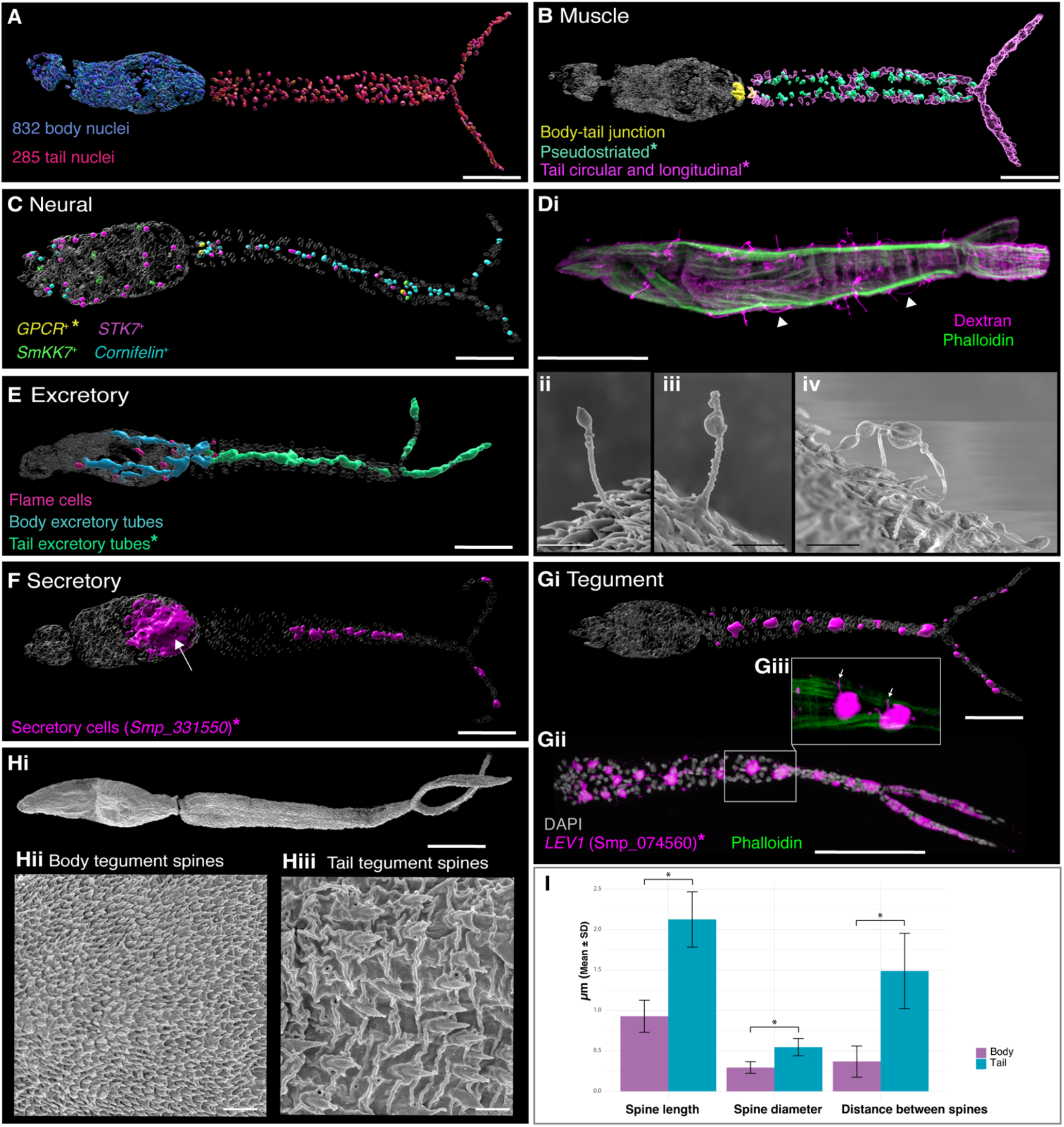
Cell type architecture of the cercarial tail. **(A)** Nuclear segmentation: 832 nuclei in the body and 285 in the tail. **(B)** 3D rendering of tail muscle marker expression. **(C)** Segmented nuclei of tail neuronal cell types. **(Di)** Dextran labelling shows cilia of tail, including ciliary loops (▼). (Dii to Div) Scanning electron micrographs (SEM) show ciliary morphologies: clubbed **(Dii and Diii)** and looped clubbed **(Div)**. **(E)** 3D rendering of excretory cell type markers expression. (F) Segmented secretory cell marker expression, also in the body acetabular glands (arrow). **(Gi)** 3D rendering of tail tegument marker, *LEV1*, expression, **(Gii)** and actual expression in large domains along the tail. (Giii) Cytoplasmic protrusions of *LEV1*+ cells (arrows) extend between muscles (phalloidin; green). **(Hi)** SEM of a cercaria, with body (Hii) and tail **(Hiii)** tegument spines magnified. **(I)** Tail tegument spines are larger and more widely spaced than body spines; *n* = 30 spines per region; Welch two-sample t tests, *P* < 0.005. Asterisks, cell types only in the tail. All scale bars, 50 μm except; Dii to Div, 1 μm; Hii and Hiii, 2 μm.

The schistosomulum atlas (Fig. 1G and fig. S1C) comprises 18,510 cells, including 13 clusters previously validated by ISH *(12)*, as well as previously unidentified clusters for excretory tubes, flame cells, and new muscle, neural and stem-like types. Between the cercarial and schistosomulum atlases, SAMap and dataset integration matched nine cercarial cell types one-to-one and four less specifically (Fig. 1H; table S4). Of the ten with no schistosomulum counterpart, six are tail-specific types.

The cercaria’s most distinctive organ—its tail—comprises ∼285 cells (Fig. 2 and fig. S11). These include the neuromuscular architecture for swimming: tail-specific smooth (circular and longitudinal) and pseudostriated muscles, as well as four neuronal types: neuropeptidergic *GPCR-B*^+^ neurons, peripheral *STK*^+^ neurons, secretory *SmKK7*^+^ neurons and ciliated *Cornifelin*^+^ neurons (Fig. 2, B and C, and figs. S4 and S5). Dextran labelling and scanning electron microscopy (SEM) revealed cilia projecting from the cercarial surface, probably corresponding to *Cornifelin*^+^ neurons. Some cilia were clubbed, whereas others, on the tail stem, interdigitated to form loops that may act as strain sensors during swimming (Fig. 2D).

We also discovered three tail-specific cell types with tentative roles beyond locomotion. A discrete excretory tube population extends towards the nephridiopores at the tips of the fork and may clear metabolic waste generated by swimming (Fig. 2E and fig. S7). A previously undescribed secretory cell type (Fig. 2F and fig. S7F) expressed secreted, putatively host-interacting genes, including *MEG-7* (Smp_165050) and *venom allergen-like 16* (Smp_124070) (data S2). Most strikingly, we found a tail-specific tegument cell type marked by *larval extracellular vesicle-1 (LEV1*, Smp_074560) (Fig. 2G and fig. S8), which encodes an immunogenic protein *(18)*. Each tail contained about 20 *LEV1+* cells, each with extensive cytoplasmic projections (Fig. 2G). The *LEV1*^*+*^ cells co-expressed the tail marker *tegument allergen-like 4 (TAL4*, Smp_169190) *(19) in silico*, and dextran-labelled cells occupied a similar position in the tail, supporting tegumental identity (fig. S8) *(20)*. This cluster also expressed further immunogens and gene products from schistosomulum extracellular vesicles (table S5) *(18, 21)*. The posterior-facing spines on the tail tegument are longer, wider, and more widely spaced than those on the body tegument (Fig. 2, H and I), suggesting they anchor the tail in the skin after penetration *(2, 4)*. There, the tail tegument and secretory cells may deposit antigens and enzymes that aid invasion or contribute to concomitant immunity. The tail is therefore not only locomotory but sensory, secretory and possibly immunogenic, with an underappreciated role in invasion.

We next asked if the proposed recent origin of cercaria is reflected at the molecular level. In SAMap comparisons, the two morphologically most distinctive tail cell types, pseudostriated muscle and tail tegument, had no clear counterpart elsewhere in the *S. mansoni* life cycle or in free-living planarian and polyclad flatworms (fig. S2; data S2), suggesting developmental and evolutionary novelty. To test whether this reflects innovation at the gene level, we assigned each *S. mansoni* gene to a phylostratum and calculated a transcriptomic age index (TAI) for each cercarial cell (Fig. 3, A and B; table S6; data S3), which is the expression-weighted mean phylostratum of a cell’s transcriptome, with higher values indicating a bias towards younger genes *(22)*.

**Fig 3.**
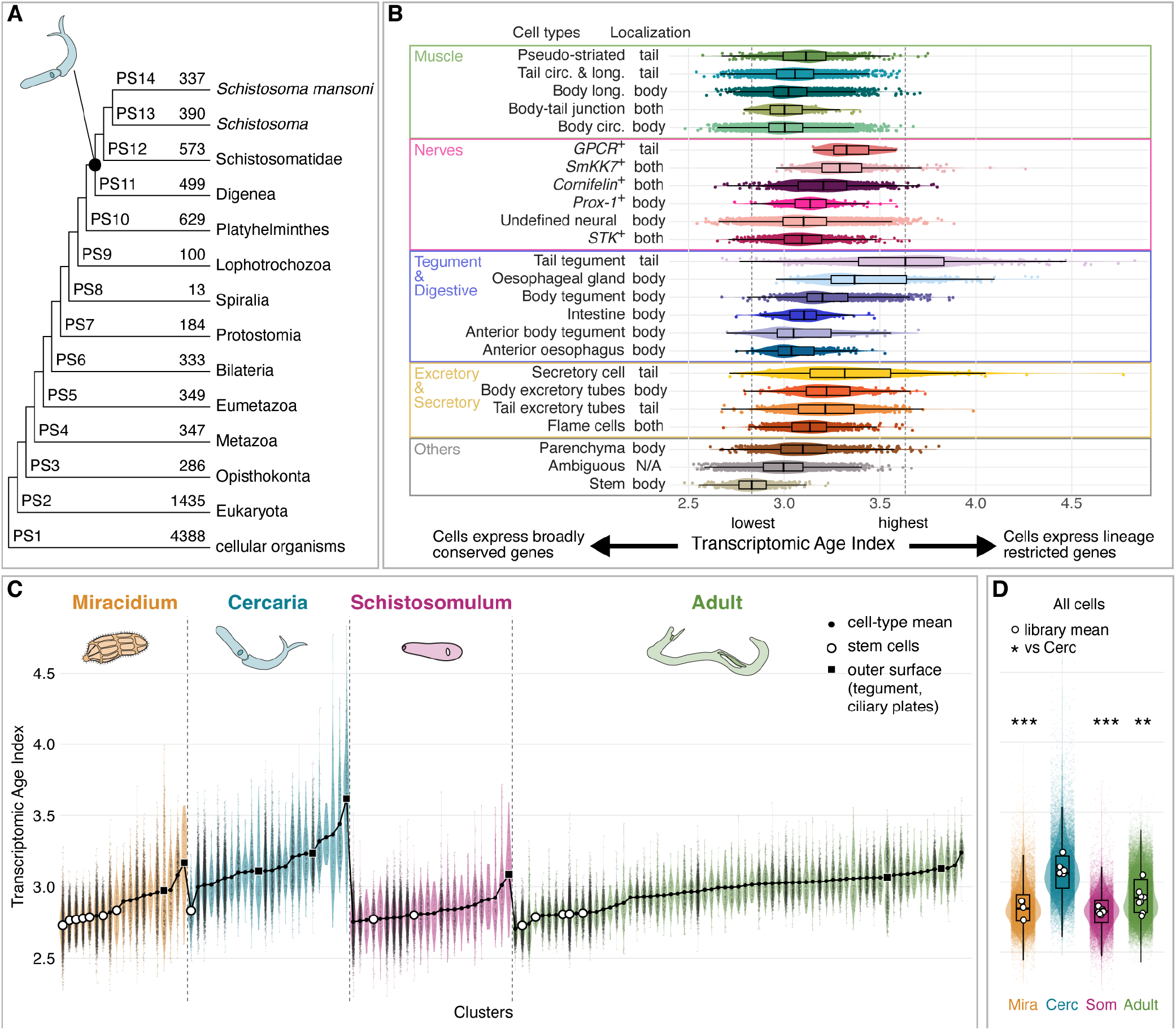
Transcriptomic age analyses of cell types. **(A)** Phylostratigraphy of *S. mansoni* genes; each gene is assigned to the oldest taxonomic level with a detectable homolog (PS1 to PS14). Numbers, genes per phylostratum; black dot, origin of digenean cercaria. **(B)** Transcriptomic age index (TAI) of cercarial cell types, grouped by tissue and TAI rank. TAI is the expression-weighted mean phylostratum of a cell’s expressed genes; higher values indicate transcriptomes shifted towards more recently evolved genes. Points, single cells; boxes, median and interquartile range (IQR); whiskers, values within 1.5 × IQR. Dashed lines, median TAI of stem cells (oldest) and tail tegument (youngest). **(C)** Single-cell TAI of each cell type in the miracidium *(14)*, cercaria, schistosomulum and adult atlases *(11)*, ordered by mean TAI within each stage. **(D)** TAI of all cells of each stage. Boxes as in (B). Cercarial TAI exceeds that of each other stage (gene-age permutation test, 10,000 permutations: *P* = 0.0007, *P* < 0.0001 and *P* = 0.004 vs. miracidium, schistosomulum and adult).

Tail-specific muscle (pseudostriated, and tail circular & longitudinal), neuronal *(GPCR-B*^+^) and tegument cell types showed higher TAI than their body counterparts (Fig. 3B). The tail tegument exhibited the highest TAI of any cercarial cell type. Because rapidly evolving genes can be assigned to younger strata, a high TAI may reflect fast sequence divergence rather than recent origin, a particular concern for genes encoding surface or secretory products. However, the tail tegument TAI remained the highest after removing genes encoding predicted secreted proteins (fig. S12; table S6), and exceeded the TAI of body tegument, which also plays a key role in the host interface. This suggests that the tail tegument expression is biased towards newer genes, rather than being ancient genes that have diverged beyond homology detection. To place cercarial cells in the context of the life cycle, we compared TAI across the cercaria, the 2-day schistosomulum and published miracidium and adult atlases (Fig. 3C). At every stage, stem cells were among the lowest-TAI populations, consistent with their ancient origin, while outer surface layers (e.g. tegument, miracidia. ciliary plates) were among the highest. The cercaria had the highest TAI of the four stages, even in cell types that persist into the schistosomulum (Fig. 3D; table S7), indicating a greater contribution of young genes to the cercarial transcriptome than to that of other stages. Together, these results provide the first cellular-level transcriptomic support for the cercaria as the most recent addition to the digenean life cycle *(6)*.

Tail cells do not appear to use a separate regulatory programme. From 398 predicted *S. mansoni* transcription factors (TFs; data S4), tail cell types expressed the same families as body cells (table S8). In cell type trees built from TFs or effector genes, each tail cell type nested within its own tissue, rather than grouping with other tail cell types (Fig. 4A; data S4). Tail identity therefore appears to be layered onto an existing programme. Each tissue shares a tissue-level TF code (in muscle, for example, *myoD, ARID3*, a *TALE homeobox* and *NFI*; Fig. 4B; table S9), many of which are retained in the same tissue at other life stages (fig. S13). Onto this, a tail-wide signature and a smaller subtype layer are superimposed. The tail-wide signature comprises two TFs with higher expression in tail than body cell types across tissues: the Hox gene *Smox1* (Smp_002640; a *fushi tarazu* orthologue) and *teashirt* (Smp_180690). Restricted to specific tail cell types are three TFs, including the Hox gene *SmHox8* (Smp_114565; an *Abdominal-A* orthologue) in tail muscles (Fig. 4C; table S8) *(23)*. Both Hox genes are central-class *(24)*. Posterior-class Hox genes were barely detected, and *teashirt* forms a posterior gradient in planarians *(25, 26)*.

**Fig 4.**
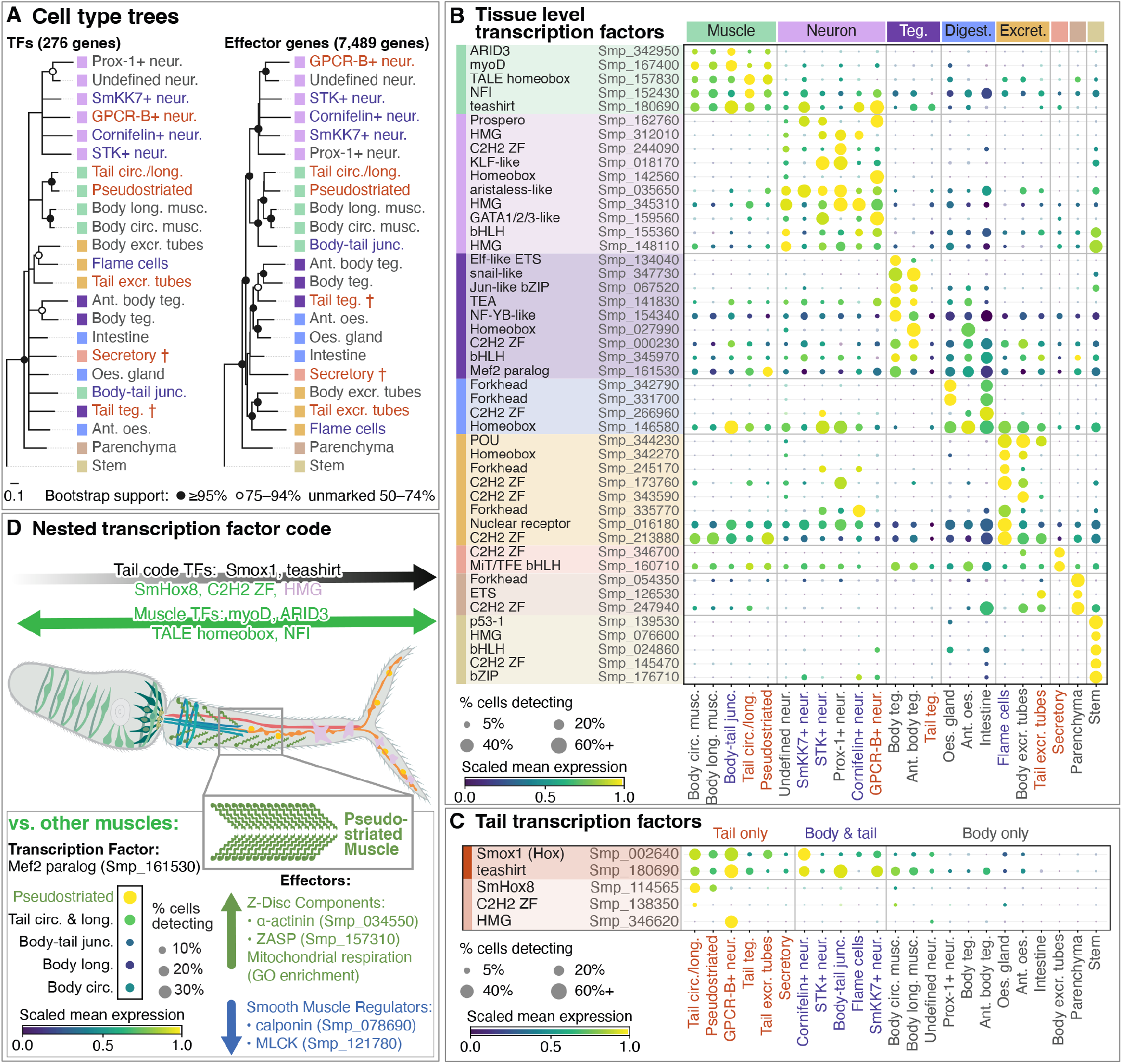
Transcription factors (TFs) in cercarial cell types. **(A)** Cell type trees from 276 TFs and 7,489 effector genes. Symbols, bootstrap support (nodes < 50% collapsed); squares, tissue; labels: red, tail only; blue, body and tail; gray, body only; †, low gene detection. **(B)** Tissue TFs: enriched against all other cells in one tissue only; shown if log_2_ fold change ≥ 1.5 and detected in ≥ 20% of cells of a member cell type (table S9). **(C)** Tail TFs (table S8). Dark bar, tail code: higher in tail than body cell types in ≥ 3 of 4 shared tissues. Light bar, restricted: enriched in a tail-only cell type, < 5% of cells elsewhere. **(D)** Nested TF codes in pseudostriated muscle, which expresses tail and muscle TFs. Versus other muscles, it has higher expression of a Mef2 paralog, Z-disc proteins α-actinin and ZASP, and respiratory chain, ATP synthesis and TCA cycle genes, and lower expression of smooth-muscle regulators calponin and myosin light-chain kinase (table S10). Tests: edgeR, pseudobulk per sex pool, > 2-fold, FDR < 0.05. Dot size, % cells detecting; color, mean expression scaled 0–1 per TF.

This nested regulatory organisation is exemplified by the pseudostriated muscle cluster, found only in the tail and with the highest TAI of any cercarial muscle type. Named for its striated-like ultrastructure *(27)* despite flatworms appearing to have lost striated muscle *(28)*, it powers tail beating during swimming. It retains the cercarial muscle TF code but expresses a *Mef2* paralogue (Smp_161530) enriched relative to other muscles (Fig. 4D). Although most of its transcriptome is shared with the **Fig**. other cercarial muscles (96.5% of tested genes are not differentially expressed [DE] vs. other muscles; data S4), the Z-disc components *α-actinin* (Smp_034550) and *ZASP* (Smp_157310) are upregulated, whereas the smooth-muscle regulators *calponin* (Smp_078690) and *myosin light-chain kinase* (Smp_121780) are downregulated (table S10). This partially mirrors smooth-to-striated transitions in other bilaterians, such as insect gut muscle *(29)*, which retain their regulatory core while gaining *ZASP* and replacing their smooth-muscle module. Flatworm smooth muscles already express the troponin-based fast module *(29)*, so here the shared changes involve the Z-disc and smooth-muscle regulators.

Furthermore, genes upregulated in pseudostriated muscle relative to other muscle populations are enriched for respiratory electron transport, ATP synthesis and TCA cycle GO terms (data S4). The tail pseudostriated muscle is thus a new cell type built from a pre-existing genetic programme: a retained muscle regulatory state with higher expression of a *Mef2* paralogue, re-weighted contractile effectors and an amplified respiratory programme. This tail-specific muscle type is sister to the other cercarial muscles and adapted for the vigorous swimming of the cercaria.

Whereas the tail and its unique cell types are discarded at transformation, the cercarial body forms the foundation of the intramammalian stage. Eleven cercarial cell types were specific to the body and five were located in both the body and tail (fig. S11; table S3). Nine had specific reciprocal counterparts in the schistosomulum: stem cells, parenchyma, flame cells, intestine, oesophageal gland, body excretory tubes, body tegument, *SmKK7*^+^ and *STK*^+^ neurons (Fig. 1H; table S4). In addition, the body–tail junction muscle showed strong alignment with the positional muscle proposed in schistosomula, which expresses *Smp_161510* along the dorsoventral midline and *wnt-2* (Smp_167140) in an anterior– posterior gradient *(12)*. In the cercaria, the junction muscle lacks *wnt-2* and expresses less *Smp_161510* but is enriched for *notum* (Smp_015820) and *teashirt* (Smp_180690) and shares enriched genes with neurons (data S2). The six junction cells of the body (Figs. 1D and 2B) may therefore persist through transformation and may be involved in patterning the positional programme of the schistosomulum.

Three body populations, however, could not be matched to a schistosomulum counterpart (Fig. 1H), suggesting that within 48h of transformation they are lost or substantially reprogrammed, or that they were not captured in the schistosomulum atlas. Two are small clusters localized anteriorly: the anterior oesophagus, which expresses markers of the adult anterior oesophagus *(30)*, and the anterior body tegument, whose position and transcriptome suggest an identity intermediate between tegument and digestive cell types associated with the head glands (data S2). The third, *Prox-1+* neurons, comprises four neurons dorsal to the brain (Fig. 1C and fig. S5; movie S1) that express GPCR signalling genes and, unexpectedly, genes for eye specification and phototransduction (data S2). They localize where other digenean cercariae have eyespots (e.g. *Lepocreadium setiferoides*; fig. S5), and the closest SAMap alignment was with planarian *opsin+* photoreceptors. Although *S. mansoni* lacks the pigmented eyes found in many flatworms, these neurons may have photoreceptive function and contribute to light-dependent responses such as cercarial shedding from snails and phototactic swimming *(31)*, which would not be required in the intramammalian stages.

Using pseudobulk differential expression, we examined how persisting tissues adapt to the intramammalian environment. Pooling the nine matched pairs into a single comparison revealed 919 genes with more than twofold higher expression in cercariae and 412 in schistosomula (Fig. 5A; data S5), and every matched pair showed the same bias (fig. S14; table S11). However, only nine genes were higher in the cercaria in all nine pairs, and only one, a heat shock protein (Smp_072330), was higher in the schistosomulum, indicating that the transcriptional change is largely cell type specific (data S5).

**Fig 5.**
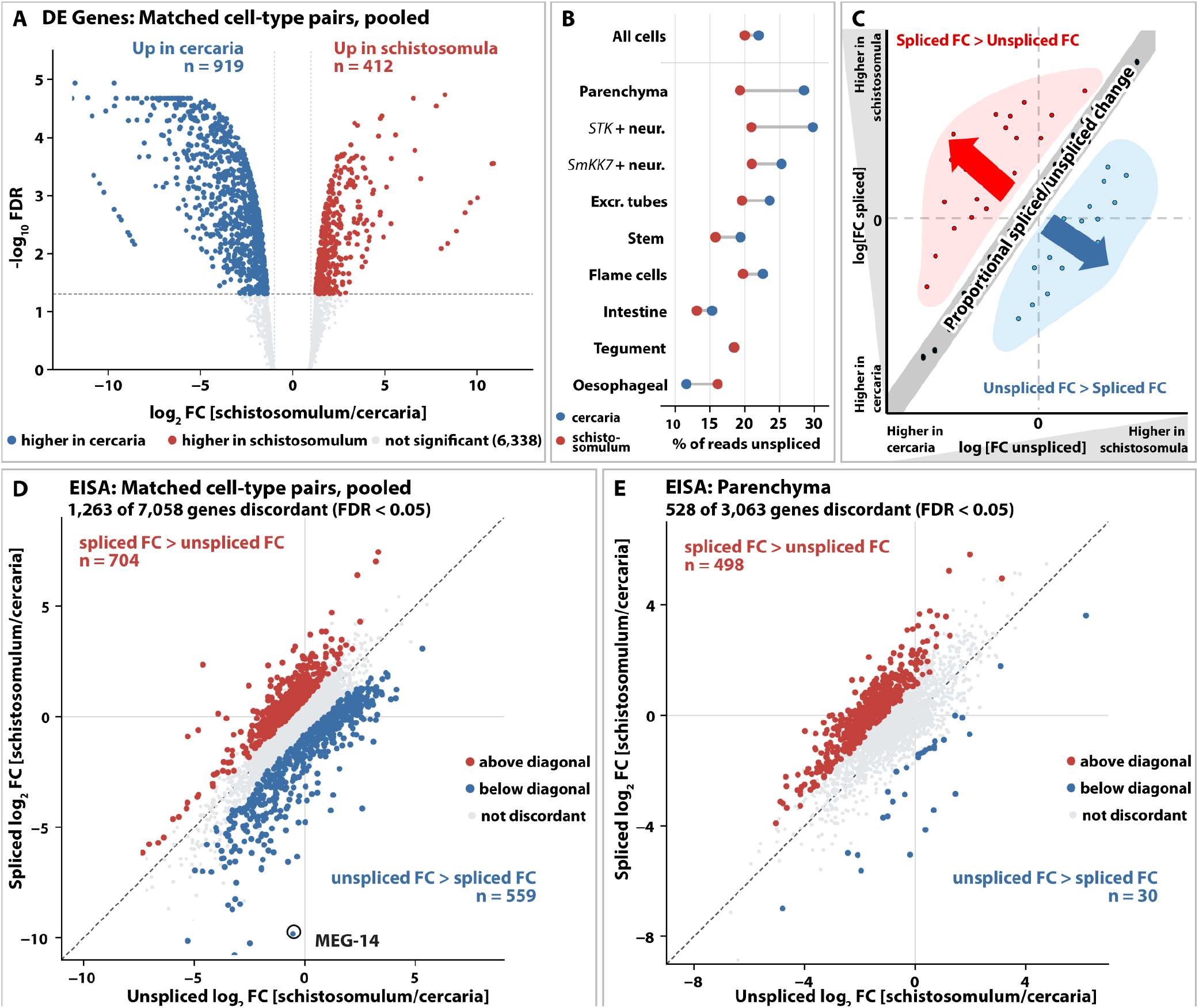
Gene expression and splicing change between matched cercarial and schistosomulum cell types. **(A)** Differential expression between cercaria and schistosomulum, nine matched cell types pooled (table S4); 7,669 genes tested. Blue, higher in the cercaria (919); red, higher in the schistosomulum (412) (edgeR, twofold threshold, FDR < 0.05, dashed line). **(B)** Percentage of reads unspliced in all cells and in each matched cell type, cercaria (blue) and schistosomulum (red); mean of the two sex pools. **(C)** Exon–intron split analysis (EISA) schematic. Genes on the diagonal change spliced (mRNA) and unspliced (pre-mRNA) reads in proportion; genes above it (red) gain more spliced than unspliced reads in the schistosomulum, genes below it (blue) the reverse. **(D and E)** EISA of the nine matched cell types pooled (D; 7,058 genes) and of parenchyma (E; 3,063 genes). Red and blue, genes significantly above (D, 704; E, 498) or below (D, 559; E, 30) the diagonal (eisaR, FDR < 0.05); *MEG-14* ringed.

An exception to this cell-type-specific reprogramming was energy metabolism. Of 30 DE genes spanning glycolysis, respiration and substrate transport, 26 were higher in the cercaria, and these were DE across more matched cell types than other DE genes (data S5). Thus, beyond the swimming tail, persisting body cells also express catabolic genes more highly than their schistosomulum counterparts. In schistosomula, two lactate dehydrogenases (Smp_038960, Smp_033040) were upregulated across multiple cell types, indicating a shift from aerobic use of stored glycogen to lactate fermentation, as in later intravascular stages *(32)*.

We observed that the transition from cercaria into schistosomulum disproportionately involves younger genes. Genes arising within the Schistosomatidae family or later were nearly three times more likely to be DE than more ancient genes (45% vs. 16%; fig. S15; table S12). Micro-exon genes (MEGs), a family of secreted, host-interacting proteins *(33)*, exemplify this: 27 of the 28 tested arose within Schistosomatidae or later, and 26 were DE in the pooled comparison, by larger factors than other DE genes of comparable age (tables S12 and S13). Their direction of expression is cell type specific. In the oesophageal gland, 18 of 20 differentially expressed MEGs were higher in cercariae. In the tegument, in contrast, *MEG-2*.*2* (Smp_345100), *MEG-2*.*3* (Smp_336990), *MEG-3* (Smp_138080) and *MEG-3*.*2* (Smp_138070) were undetectable in cercariae yet among the most abundant schistosomulum transcripts, whereas a *MEG-14* (Smp_330180) was among the most abundant cercarial tegument transcripts but nearly absent in the schistosomulum (fig. S15; table S13). This suggests that the oesophageal gland reduces its cercarial MEG programme across the transition while the tegument switches to a new set.

To investigate how cercaria-biased transcripts are depleted and new transcripts are made in the schistosomulum, we classified the reads in both atlases as either unspliced (intron-containing) or spliced (exon-only). Unspliced RNA is conventionally interpreted as a proxy for active transcription and spliced RNA as mature mRNAs. If cercariae are transcriptionally quiescent *(10)*, we hypothesised that their unspliced fraction should be lower than in the actively remodelling schistosomulum.

Unexpectedly, unspliced transcripts comprised a similar proportion of reads in both stages (22% and 20% across all cells in cercariae and schistosomula respectively; Fig. 5B). These proportions of reads varied among matched cell types: from the highest cercarial excess in parenchyma (28.5% vs. 19.3%) to the reverse in oesophageal gland (11.6% vs. 16.1%). This suggests that, if there is quiescence, it is incomplete and variable, or that the pre-mRNA signal is due to storage of pre-mRNA *(8)* or intron retention *(34)*.

RNA velocity models fit very few genes across the transition (fig. S16), as would be expected if cercariae truly are quiescent. We used exon-intron split analysis (EISA) *(35)*, which compares changes in unspliced reads with spliced reads for each gene. Discordant changes are usually interpreted as changes after transcription, such as altered mRNA stability or processing (Fig. 5C). Across the nine matched cell types, 1,263 of 7,058 genes (17.9%) were discordant (FDR < 0.05; Fig. 5D; data S6). If cercariae carry a stored pool of mature transcripts cleared after transformation, most discordant genes should lie below the diagonal of a spliced vs. unspliced log[fold-change] plot (Fig. 5C). Instead, both directions were common (704 above, 559 below). Some below-diagonal genes, such as *MEG-14* (Smp_330180), showed large losses of spliced transcripts with little change in unspliced, consistent with clearance of stored mRNA. Discordance also differed between cell types (fig. S17); in parenchyma 94% of discordant genes lay above the diagonal (Fig. 5E; data S6), mostly because unspliced reads fell more than spliced reads. These above-diagonal genes carried approximately 1.5-fold more unspliced RNA in cercariae compared with other parenchyma genes taking into account expression and intron length (1.25- and 1.73-fold in two independent halves of the data). We have two interpretations for these observations. If cercarial parenchyma is quiescent, unspliced reads must represent stored unprocessed RNA that is depleted after transformation, with the enrichment suggesting both tissue- and gene-specific storage differences. Another possibility is that parenchyma is transcribing and the shift simply reflects lower transcription after transformation or differences in splicing kinetics. This is feasible given that a small transcriptional signal was previously observed in cercariae, and it was assessed over 18h, long enough for early-labelled RNA to have decayed *(10, 36)*. With either scenario, our cercarial parenchyma finding challenges the stage-wide near-total quiescence model and shows that the transcriptional dynamics of the transition are more complex than previously reported, with variability between tissues. In at least some tissues, cercariae are either still transcribing 4 hours after emergence from the snail, or a significant proportion of any stored RNA is unprocessed pre-mRNA, which has previously not been considered.

There is an increasing number of single-cell atlases of schistosomes but important gaps in knowledge persist. In this paper we present a systematic atlas of the human-infective cercaria and a complementary atlas tracking the transition in the intramammalian schistosomulum. Together, these atlases show that the cercaria has cellular novelty as well as being morphologically distinct: its transient tail comprises cell types found nowhere else in the life cycle or in other flatworms, built by layering a tail-wide regulatory signature onto existing tissue programmes and drawing disproportionately on young genes. That tail-specific cell types are the youngest in the cercaria, and cercarial cells the youngest yet detected across the life cycle, suggests the shift from ingestion-based to tail-driven active transmission was built on cellular innovation and specialization within tissues allowing dispersal and the chance to infect more than one definitive host (fig. S18). We uncovered complex transcriptional dynamics associated with the transformation, and distinguishing stored transcripts from ongoing transcription will be a key next step in understanding how this short-lived larva prepares for transformation in the human host. Many body cell types persist into the schistosomulum, modify their energy metabolism, and undergo cell-type-specific reprogramming, including the swap of micro-exon gene repertoires. Our findings highlight the immunogenic products of the tail tegument and the genes turned over at transformation as candidate targets for blocking host invasion.

## Supporting information

Movie S1. Prox-1+ and SmKK7+ neurons in the cercarial body.

Data_S2_cell_type_markers_and_matching

Data_S3_gene_ages

Data_S4_transcription_factors

Data_S5_cercaria_schistosomulum_DE

Data_S6_EISA

## ACKNOWLEDGEMENTS

We thank Karl Hoffmann and his team (Aberystwyth University) for providing *Schistosoma mansoni*-infected mouse livers from which we extracted eggs for mono-miracidial infections of *Biomphalaria glabrata* snails to collect single sex cercariae and schistosomula. We thank Simon Clare, Catherine McCarthy and Lisa Seymour for technical assistance with the life cycle maintenance and collection of parasite material at the Wellcome Sanger Institute. We thank the Schistosomiasis Resource Center of the Biomedical Research Institute (Rockville, MD), which receives support from National Institute Of Allergy And Infectious Diseases of the NIH (R24AI19813), for sending exposed *Biomphalaria glabrata* snails for validation of cercarial cell type experiments. FACS and RNA-seq were performed with the aid of the Wellcome Sanger core facilities. We thank Carsten Wolff and Louis Kerr from the Core Microscopy Facility at the MBL for technical assistance with image processing in Imaris and SEM sample preparation and imaging. We thank Ana Corum for the collection and imaging of *Lepocreadium setiferoides*.

## Author contributions

Conceptualization: M.B., G.R., K.R., T.A.; Data curation: T.A., S.O.K., Y.X., K.R., F.B., G.R.; Formal analysis: T.A., Y.X., F.B., S.O.K.; Funding acquisition: K.R., M.B.; Investigation: S.O.K., Y.X., M.L., J.G., S.K., T.A., G.R., K.R.; Project administration: K.R., M.B.; Resources: M.B., K.R., G.R., M.L.; Supervision: G.R., M.B., K.R.; Validation: KR, TA; Visualization: S.O.K., Y.X., T.A., K.R.; Writing - original draft: S.O.K., Y.X., T.A., K.R., M.B.; Writing – review and editing: Y.X., G.R., F.B., T.A., M.B., K.R.; Designing web-based resources: Y.X.

