## Supplementary material for "Cell atlases of schistosome human infective stages reveal larval tissue fates and innovations": Methods, Supplementary Text and Figures

### Materials and Methods

#### Ethics statement

At the Wellcome Sanger Institute (WSI), the life cycle of *Schistosoma mansoni* (NMRI strain) was maintained by infecting in-house bred freshwater *Biomphalaria glabrata* snails (NMRI strain) and outsourced mice (outbred TO). All regulated procedures were performed under Home Office Project Licence No. P77E8A062 held by GR, approved by the Animal Welfare and Ethical Review Body of the WSI, as required by the UK Animals (Scientific Procedures) Act 1986 Amendment Regulations 2012.

#### Parasite material and cell dissociation

Cercariae were collected from snails infected with single miracidia, and their sex was determined as described (*37*): 4 to 7 weeks after exposure, individual snails were placed in 12-well plates with 4 ml of water per well and left to shed cercariae under light for ~4 hours, and cercariae from each snail were concentrated by centrifugation, lysed and typed by end-point PCR with a W chromosome-specific primer set (*38*) and an *actin-2* (Smp_307020, in the pseudoautosomal region of the Z chromosome) primer set (*39*). Snails were then kept in 'male' and 'female' tanks, from which male or female cercariae and schistosomula were collected. Schistosomula were produced by mechanical transformation of male or female cercariae and cultured for 48 hours (*12*).

For each stage, one female and one male sample was collected (cercaria: ~42,500 female and ~27,000 male; 2-day schistosomulum: 20,000 female and 22,000 male); cercariae and schistosomula were processed on different days with the same protocol. Larvae were filtered through a 100 µm filter, concentrated on ice for 1 hour in Low Protein Binding 15 ml tubes, and all but 20 µl of water (or schistosomulum wash medium) was removed. Larvae were digested in 300 µl of digestion buffer (2.5 µl of 0.5 M EDTA in 300 µl of HBSS without $Ca^{2+}$ and $Mg^{2+}$) with 200 µl of Native *Bacillus licheniformis* Protease (NATE-0633, Creative Enzymes; 100 mg/ml stock, 40 mg/ml final) (*40*) for 70 minutes on ice, pipetting gently ~20 times every 5 minutes. The enzyme was quenched with 200 µl of 25% BSA in HBSS, diluted with 10 ml of ice-cold PBS (with $Ca^{2+}$ and $Mg^{2+}$) and the cells centrifuged at 350 g for 5 minutes at 4 °C. Cells were then digested with 100 µl of Liberase TM (Roche; 2.5 mg/ml stock, 2.08 mg/ml final) for 20 minutes at room temperature, pipetting ~10 times every 5 minutes, passed through a 30 µm strainer, resuspended in 5 ml of cold PBS with 0.5% BSA, centrifuged at 350 g for 10 minutes at 4 °C and concentrated to 1,200 µl. From each sample, 500 µl was stained with propidium iodide (1 µg/ml; Sigma P4864) and live (PI-negative) cells were sorted on a BD Influx cell sorter, with unstained cells as an autofluorescence control; 500 µl was left unsorted to preserve cell diversity. This gave one sorted and one unsorted suspension per sex and stage.

#### Single-cell library preparation and sequencing

Libraries were prepared with Chromium Next GEM Single Cell 3′ Reagent Kits v3.1 (10x Genomics; user guide CG000204 Rev D). Cell concentrations were estimated by flow cytometry (Beckman Coulter CytoFLEX S) and adjusted appropriately. Two aliquots of each suspension were loaded, targeting ~10,000 cells per reaction, giving eight reactions (one chip) per stage (fig. S1B; table S1). The eight libraries of each stage were sequenced on one lane of an Illumina NovaSeq (XP workflow; read 1, 28 bp; read 2, 90 bp; index reads, 10 bp each)..

### Read mapping and quantification

Reads were mapped to the *S. mansoni* v10 genome (*41*) from WormBase ParaSite (*42*) with Cell Ranger 7.0.1 (*43*), excluding introns, after modifying *Smp_323380* and *Smp_337410* as described (*14*) to reduce misassignment of reads between Z and W gametologues; read quality was checked with FastQC 0.11.8 (*44*). In the cercaria, 47.99% of reads mapped confidently to the transcriptome (44.2–53.5% per library), and in the schistosomulum 61.75% (58.1–69.1%). The eight cercarial libraries yielded 30,158 cells (sorted: 2,413 male, 3,259 female; unsorted: 6,694 male, 17,792 female), with a median of 1,019 UMIs and 592 genes per cell (table S1); the eight schistosomulum libraries yielded 19,857 cells (sorted: 3,540 male, 4,651 female; unsorted: 5,518 male, 6,148 female), with a median of 1,237 UMIs and 824 genes per cell (medians of the per-library medians).

### Quality control and ambient RNA

Data were analysed in R 4.4.1 with Seurat 5.1.0 (*45*). Cells with < 50 genes and genes detected in < 3 cells were removed, and doublets were predicted with scDblFinder 1.18.0 (*46*). Thresholds were chosen from the distributions of mitochondrial and ribosomal read fractions, UMI and gene counts and doublet calls across the eight libraries after exhaustive initial analyses: cells with < 5% mitochondrial reads, > 500 UMIs, > 140 genes and a singlet call were retained, without a ribosomal filter, to remove damaged cells while keeping unknown cell types. A stricter mitochondrial threshold was not used because the two clusters with the highest mitochondrial fraction had clear marker genes. After QC, 24,708 cercarial and 18,510 schistosomulum cells remained (schistosomulum filters were established in the same manner: < 5% mitochondrial reads, > 500 UMIs, > 167 genes and a singlet call were retained). Ambient RNA was estimated per library with SoupX (*47*) before QC filtering (SCTransform (*48*), 30 PCs, default parameters, no predefined marker genes). The estimated contamination fraction for the cercarial samples was 1–8% and 1-3% for the schistosomula, so counts were not corrected; the top predicted cercarial contaminants (*Smp_033000*, *Smp_003770* and *Smp_163710*, in all eight cercarial libraries) were considered when interpreting genes of interest.

### Clustering, cell-type annotation and integration

The data for both were processed following the standard Seurat workflows: the data were normalised, variable features selected (n = 2000), PCs were calculated for n = 100. The data were also prepared using SCTransform. The final number of PCs, neighbours, and clustering resolution was calculated using JackStraw, Elbow plots, and Silhouette scores for cercariae (PCs = 26, resolution = 0.6), and for schistosomula (PCs = 25, resolution = 0.4). For both atlases, cell-type annotations were made using expression of previously validated markers and top cluster genes. The atlases were integrated using anchors on the SCTransformed data, which were reclustered together (PCs = 36, resolution = 0.5).

Gene annotations from the v10 genome were used; descriptions of marker genes were inspected manually and revised using InterPro and PANTHER families where necessary. Marker genes of each cell type were identified with Seurat FindAllMarkers (ROC test) (table S2; data S2).

### *In situ* hybridization, immunofluorescence and fluorescent labelling

Whole-mount hybridization chain reaction (HCR v3) (*49*) was performed on cercariae for 22 cell-type marker genes and four further genes, with probes, buffers and hairpins from Molecular Instruments designed against WormBase ParaSite WBPS19 sequences (table S3). Cercariae were

shed into pond water under light, collected in 35 μm incubation baskets (Intavis) in 24-well plates and, within 2 hours, fixed in 4% paraformaldehyde in PBSTw (DEPC-treated, 0.22 μm-filtered PBS with 0.1% Tween 20) for 30 minutes at room temperature, washed, permeabilized with 60 μg/ml Proteinase K (Invitrogen) for 25 minutes at 37 °C, then incubated in 2 mg/ml glycine in PBSTw on ice for 15 minutes, before washing in PBSTw. Samples were pre-hybridized in 1:1 PBSTw:hybridization buffer for 5 minutes and in hybridization buffer for 1 hour at 37 °C, hybridized overnight at 37 °C with 2.6 nM of each probe, and washed four times for 15 minutes in probe wash buffer at 37 °C and three times for 5 minutes in 5× SSCTw. After 30 minutes in amplification buffer, samples were incubated overnight in the dark at room temperature with snap-cooled hairpins (95 °C for 90 seconds, 30 minutes at room temperature; 6 nM final), and washed four times in 5× SSCTw.
Some cercariae were then immunolabelled: they were blocked for 1 hour (5% horse serum, 0.45% fish serum, 0.05% Tween 20 in 5× SSC with 0.3% Triton X-100 [5× SSCTx]), incubated overnight at 4 °C with primary antibodies against acetylated tubulin (Sigma [T7461], 1:500), phosphotyrosine (Cell Signaling Technology [P-Tyr-100], 1:100) or a custom antibody against the neuropeptide SmNPP-35 (Smp_188580; Pocono Rabbit Farm, 1:100), washed for 1 hour, incubated overnight at 4 °C with secondary antibodies [Invitrogen; goat anti-mouse 488 [A28175], goat anti-rabbit 633 [35562], 1:500] and washed for 2 hours. Most cercariae were labelled with Alexa Fluor 488 phalloidin (Invitrogen; 1.5:500) for 5 hours at room temperature. Gland cells were labelled with Cy5-conjugated peanut agglutinin (Vector Laboratories; 1:500, overnight at 4 °C) (*50*). Cercariae were mounted in DAPI Fluoromount-G (SouthernBiotech) and imaged on a Leica SP8 confocal microscope. Each marker gene was tested in more than three experiments, imaging > 10 cercariae per experiment. HCR controls to test for hairpin trapping and non-specific probe binding were carried out, and all immunofluorescence experiments included no-primary antibody and no-primary/no-secondary antibody controls, and were imaged at identical settings, to exclude background and autofluorescence.

**Tail tegument labelling**

Tail tegument cell bodies were labelled with fluorophore-conjugated dextran (D3312, Invitrogen) as adapted from (*20*). Freshly shed cercariae were collected in a 35 μm basket, agitated in 5 mg/ml dextran in nuclease-free water for 5 minutes by pipetting, fixed in 4% paraformaldehyde in PBSTx (PBS with 0.3% Triton X-100) for 30 minutes, blocked in PBSTx with 1% BSA for 1 hour, labelled with phalloidin (1.5:500, 4 hours) and mounted in DAPI Fluoromount-G for confocal imaging. Cercariae that had lost their tails mechanically showed the best tegument labelling; dextran also labelled cilia and ciliated cell bodies.

**Image segmentation and nuclear counts**

Confocal z-stacks were segmented in Imaris (Oxford Instruments, v10.0.0). Nuclei were counted with the 'spots' function (table S3) and segmented for display with the machine-learning and manual 'surfaces' functions, and staining signal was rendered with intensity-based 'surfaces'. Nuclei most closely associated with marker gene signal were assigned to that cell type. To compare more than three genes or labels (e.g. the five muscle markers, or PNA staining with the secretory marker), z-stacks from different samples were aligned on the DAPI signal and overlaid.

**Scanning electron microscopy**

Cercariae were shed and collected in 35 µm incubation baskets in 24-well plates, fixed in 4% paraformaldehyde and 0.5% glutaraldehyde in PBS for 1 hour at room temperature, rinsed four times in PBS and dehydrated on ice through a graded ethanol series in PBS (50%, 70%, 85% and 95%, 10 minutes each; 100%, three times for 15 minutes). Samples were critical-point dried with $CO_2$ (Leica CPD300) in 30 µm specimen baskets, transferred individually with a metal probe onto stubs with carbon adhesive tabs, sputter-coated with 4 nm of platinum and imaged on a Thermo Fisher Helios 5 Hydra PFIB-SEM at 3 kV with the through-lens detector. Tegument spines were measured in Fiji: length, diameter, and distance to the nearest neighbouring spine, for 30 spines each in the body and tail (Fig. 2I; data S1). Body and tail were compared with Welch two-sample *t* tests.

**SAMap alignment**

Cell types were aligned across life stages and species with SAMap 3.0.1 (*17*). Raw count matrices of the quality-controlled atlases, annotated with their cell types, were imported as AnnData objects: cercaria and 2-day schistosomulum (this study), miracidium (*14*), adult (*11*), the planarian *Schmidtea mediterranea* (*15*) and the polyclad *Prostheceraeus crozieri* (*16*). Gene homology graphs were built from reciprocal BLAST searches with SAMap's map_genes.sh (E-value $\leq 10^{-6}$, one high-scoring pair per hit); between *S. mansoni* stages, which share a genome, this reduces to identical genes, so the alignment rests on the cell–cell mapping alone. SAMap was run with default parameters and the annotated cell types as keys in two joint analyses, one of the four *S. mansoni* stages and one of the cercaria with the two free-living flatworms, so that within- and between-species comparisons did not share a manifold; cross-dataset neighbourhoods were computed for each pair of datasets (run(pairwise=True)), and alignment scores between cell types were computed with get_mapping_scores (data S2).

**Matching cercarial and schistosomulum cell types**

Cercarial and schistosomulum cell types were matched by combining four lines of evidence: localization by HCR, SAMap alignment (above), the 50 nearest neighbours of each cell in the first 36 principal components of the integrated cercaria–schistosomulum dataset, and shared marker genes. On this evidence each cercarial cell type was classed as a specific reciprocal match, matched but not specific, or cercaria-only (table S4). The nine specific reciprocal matches were used for the comparisons below.

**Gene ages and transcriptome age index**

Gene ages were assigned with GenEra version 1.4.1 (*51*) for the WBPS19 *S. mansoni* proteome (PRJEA36577) against the NCBI non-redundant protein database (nr) and taxonomy dump (downloaded on 27 October 2025; E-value 1e-5), placing each gene in the oldest of 14 phylostrata in which a homolog was detected and taking one dated isoform per gene (9,863 dated genes; table S6; data S3). Signal peptides were predicted with SignalP 6.0 *(52)* (eukarya; Sec/SPI probability $\geq 0.5$) and subcellular location with DeepLoc 2.1 (*53*); proteins with a signal peptide or an extracellular prediction were called secreted. The transcriptome age index (TAI) *(22)* of each cell, the mean phylostratum of its dated genes weighted by log-normalized expression ($\log(1 + \text{counts per } 10^4)$; Seurat NormalizeData), was computed with myTAI 2.3.5 (*54*) for the cercaria, schistosomulum, miracidium (*14*) and adult (*11*) atlases. Differences in mean TAI between atlases were tested by permuting phylostrata among dated genes (10,000 permutations; two-sided), and between matched cercarial and schistosomulum cell types with an exact two-

sided Wilcoxon signed-rank test (SciPy 1.17) *(55)*. TAI was recomputed after removing genes predicted to encode secreted proteins: signal peptide (715 genes), extracellular (555), both (363) or either (897), counting dated genes present in either atlas (data S3). The removed genes were left out of the TAI but not out of the normalization (fig. S12; table S6).
The TAI results were further evaluated by removing phylostrata one-by-one (phylostrata 6 to 14), by removing the genes indicated as possible contaminants by SoupX, and by removing the full set of MEGs. None of these checks resulted in major changes, and the finding of cercarial tegument as the youngest cluster, and of cercarial cell clusters as younger than schistosomulum clusters persisted across all these evaluations.

**Transcription factor catalogue**
The WBPS19 proteome was searched against Pfam-A 38.2 (*56*) with hmmsearch (HMMER 3.4) (*57*) at gathering thresholds (--cut_ga). Proteins with a DNA-binding domain (DBD) were retained, excluding those with only weak or transposon-associated DNA-binding motifs (AT-hook, BED, CENP-B, psq) or with a transposase, integrase or reverse-transcriptase domain. Proteins without a DBD were rescanned against the DBD models alone (hmmsearch -T 3 --domT 3) and added if they had an HLH domain (≥ 10 bits) or ≥ 3 C2H2 zinc fingers (≥ 3 bits, one ≥ 10 bits). TFs annotated by KEGG, GO or CIS-BP v3.10 (*58*) whose DBD was missing from the current gene model were added after manual review, giving 398 TFs (data S4). Families of TFs named in the text were assigned with maximum-likelihood trees of the DBD (± 10 residues; ± 35–40 residues for bHLH-zip and MADS domains) that included *S. mansoni* paralogs, the top BLASTP hits in the planarian *Schmidtea mediterranea* (dd_Smed_v4) *(59)* and the polyclad *Prostheceraeus crozieri* (*60*), and reviewed human and *Drosophila* UniProt proteins. Domains were aligned with MAFFT v7.526 (*61*) (L-INS-i: --localpair --maxiterate 1000) and trimmed with trimAl v1.5 (*62*) (-gt 0.2); Teashirt, Snail and NFI were aligned as full-length proteins and trimmed with trimAl -automated1. Trees were inferred with IQ-TREE 3.1.4 (*63*) with ModelFinder model selection (*64*) (for nuclear receptors, limited to Q.INSECT, Q.PFAM, LG, JTT and WAG) and 1,000 ultrafast bootstraps (*65*). Specific names were used only when supported by these trees or a published *S. mansoni* name, and family-level ("-like") or domain-level names otherwise.

**Gene enrichment in cell types**
Raw counts were summed per cell type and sex pool (pseudobulk), excluding the unassigned cluster. Because the eight libraries of each stage are sorted and unsorted aliquots of one male and one female sample, the two sex pools were the replicates; pools with < 5 cells of the cell type were dropped, and a test was run only when both remained. Genes were filtered with filterByExpr (min.count = 3, min.total.count = 10), normalized by TMM and fitted with quasi-likelihood negative binomial models in edgeR 4.10 (*66*) (R 4.6; glmQLFit, design ~ sex pool + group). Every enrichment and differential expression test in this study was made against a twofold threshold (glmTreat, lfc = 1) (*67*) with a Benjamini–Hochberg FDR < 0.05. Each cell type was compared with all other cells and with the other cell types of its tissue; genes passing the test and detected in ≥ 5% of the cell type's cells were called enriched (data S2; TFs in data S4). The within-tissue test of pseudostriated muscle against the other four muscle types is in table S10 and data S4.

**Gene Ontology enrichment**

Direct GO annotations were taken from WormBase ParaSite BioMart (WBPS19) (*42*). Each set of enriched genes defined above, and the genes up and down in pseudostriated muscle, was tested for over-representation of terms in each of the three GO domains with Fisher's exact test as implemented in topGO 2.64 (*68*) (classic algorithm, nodeSize = 5), against the genes tested in the same contrast carrying ≥ 1 GO term (6,644 of 8,699 for the cell-type tests; 5,913 of 7,544 for pseudostriated muscle). *P* values were corrected within each gene set and GO domain (Benjamini–Hochberg), and terms with FDR < 0.05 are reported with their odds ratios (data S2 and data S4). The same procedure was used for every GO test in this study. Translation and ribosome terms recur in the cell types with the highest ribosomal protein content and were not interpreted.

**Transcription factor codes**

37 TFs enriched in tail-only and 122 body-only (undefined neural counted as body) cell types (table S8) were compared by family with Fisher's exact tests (Benjamini–Hochberg correction). For the tail code, tail and body cell types were compared within each of four tissues found in both regions (tegument, excretory tubes, muscle and neurons), so that tail TFs could be distinguished from tissue TFs. All four comparisons were fitted in one edgeR model, with one group per tissue and location and a tail-versus-body contrast within each tissue. Only cell types restricted to the tail or to the body were used (e.g. for neurons, *GPCR-B*$^+$ versus *Prox-1*$^+$ and undefined neural). TFs enriched in the tail in ≥ 3 of the 4 tissues formed the tail code (Fig. 4C; table S8). TFs enriched in a tail-only cell type and detected in < 5% of cells of every other cell type were called restricted to tail cell types (Fig. 4C). For the tissue code, the cell types of each tissue with several cell types (muscle, neurons, tegument, digestive, excretory) were pooled and compared with all other cells, requiring higher expression in at least two-thirds of them and detection in ≥ 5% of cells of one; secretory cells, parenchyma and stem cells, each a single cell type, used the cell-type test. TFs enriched in only one of these eight tissues were called tissue-specific; Fig. 4B shows those with log2FC ≥ 1.5 detected in ≥ 20% of cells of a member cell type (all results in table S9).

**TF use across life stages**

The cell-type and tissue tests were repeated on the miracidium (*14*), 2-day schistosomulum and adult atlases (*11*), with the two sex pools as replicates for the schistosomulum and the libraries for the miracidium (four libraries, sex not recorded) and adult (nine libraries: three each of males, mature females and immature females), which lack comparable sex pools. For TFs enriched in tail-only cell types, we recorded enrichment in other cercarial and other-stage cell types (table S8). The tissue TFs of Fig. 4B were tested in each atlas with the same test, with cell types grouped by tissue according to that atlas's own annotation: a TF was enriched in a tissue if the tissue's cell types pooled against all other cells (or, for a tissue of one cell type, that cell type against all other cells) gave FDR < 0.05 against the twofold threshold and the TF was detected in ≥ 5% of cells of a member cell type. The Fig. 4B requirement that the TF be higher in two-thirds of the tissue's cell types was not applied, because atlases divide tissues into different numbers of cell types. The number expected by chance in each TF's own tissue was the share of expressed TFs (≥ 5% of cells of a cell type of that atlas) enriched in that tissue, summed over the testable TFs (fig. S13).

**Cell type trees**

Each cell type was represented by its pseudobulk expression (log(CPM + 1)) of the 276 TFs or 7,489 other genes detected in ≥ 5% of cells of any cell type. Pairwise distances were 1 − Spearman's ρ over genes detected in ≥ 10% of cells of either cell type, and neighbour-joining trees were built with Biopython 1.88 (*69*) (DistanceTreeConstructor) and rooted on stem cells. Node support was the percentage of 1,000 trees built from genes resampled with replacement; nodes with < 50% support were collapsed (Fig. 4A). Trees built with 5% or 20% detection cutoffs or with Euclidean distance did not group the tail cell types, apart from tail tegument with secretory cells, both with low gene detection (data S4).

**Cercaria–schistosomulum differential expression**

UMI counts were summed per stage × sex pool × cell type, with the two sex pools as replicates; pseudobulks of < 10 cells were dropped. Genes were filtered with filterByExpr and then aveLogCPM > 2, normalized by TMM and fitted with edgeR quasi-likelihood models with the sex pool as a blocking factor, and DE was tested with glmTreat (lfc = 1; FDR < 0.05) for each matched pair and for the nine pairs pooled (data S5; table S11). Using the 16 libraries as units gave the same fold changes (r = 0.97–0.997). Energy-metabolism genes were assigned by matching WBPS19 gene descriptions to 46 enzyme steps (data S5), and the number of pairs in which each pooled DEG was also DE was compared between energy and other DEGs (two-sided Mann–Whitney test). Young (phylostratum ≥ 12, Schistosomatidae or younger) and older genes were compared for DE rate (Fisher's exact test) and for |log2 fold change| in each direction (table S12), as were young DE MEGs (table S13) and other young DE genes (two-sided Mann–Whitney tests; SciPy 1.17) (*55*).

**Spliced and unspliced counts and RNA velocity**

Reads of the eight cercarial and eight schistosomulum libraries were pseudoaligned to the *S. mansoni* v10 genome (WBPS18, with the same gametologue modifications) with the lamanno workflow of kallisto bustools 0.26.4 (*70*, *71*) to obtain spliced and unspliced counts. Matrices were filtered with DropletUtils 1.24.0 (*72*) and BUSpaRse 1.14.1 (*73*) and restricted to cells that passed QC in Seurat. The unspliced fraction is the unspliced share of all spliced and unspliced reads, computed for all cells and for each matched cell type (mean of the two sex pools). For RNA velocity, AnnData objects were built with Scanpy 1.11.5 (*74*) (Python 3.14.2), filtered as in Seurat (≥ 140 genes and ≥ 500 counts per cell; genes in ≥ 3 cells), normalized with scVelo 0.3.3 (*75*) (filter_and_normalize, min_shared_counts = 20, top 2,000 genes) and analysed with the dynamical model with default parameters. For matched cell types, top dynamical genes expressed mainly in those cell types and differentially expressed between stages were shortlisted, and their phase portraits were judged for fit to the model by a colleague blind to their identity.

**Exon–intron split analysis**

Spliced and unspliced counts (above) were pseudobulked per stage × sex pool × matched cell type as for DE and analyzed with eisaR 1.22.1 (*35*), *(76)*, with the two sex pools as replicates, for the nine matched cell types pooled and for each pair (data S6). Genes whose spliced and unspliced changes differed (FDR < 0.05) were discordant: below the diagonal when spliced RNA changed less than unspliced RNA from cercaria to schistosomulum, above it otherwise.

### Supplementary Text

#### Descriptions of cercarial cell types

#### S1. Muscle

Phalloidin staining of filamentous actin with ISH of the pan-muscle marker *myosin heavy chain II* (*MHC II*, Smp_085540) resolved a meshwork of circular, longitudinal and diagonal fibres in the body wall and the muscular tail (fig. S4, A and B). Searching the atlas for *myosin light chain* (Smp_045220), *troponin T* (Smp_018250), *myosin regulatory light chain* (Smp_132670) and *tropomyosin-1* (Smp_340760) identified five muscle clusters.

**S1.1 Body circular and acetabular anchor muscles** (6,157 cells)
Top marker gene *cysteine dioxygenase* (*CDO*, Smp_165750) was detected in 25% of cells of the cluster and localised around ~50 nuclei of the circular body-wall muscles and the muscles anchoring the acetabulum to the body wall (Fig. 1 and fig. S4D). Circular body-wall nuclei aligned with the dorsal and ventral midlines.

**S1.2 Body longitudinal muscles** (2,097 cells)
*Uroporphyrinogen-III synthase* (*URO3S*, Smp_079840) was detected in 46% of cells and localised around ~100 nuclei in five bands encircling the body, matching the five interdigitating tiers of longitudinal body-wall fibres (Fig. 1; fig. S4D).

**S1.3 Body–tail junction muscles** (127 cells)
*E3 ubiquitin-protein ligase* (Smp_332470) was detected in 95% of cells and localised to a ring of six nuclei at the posterior of the body and two at the anterior of the tail, with weaker signal around the brain (Fig. 1; fig. S4E). Other top markers, *gelsolin* (Smp_008660), which severs actin filaments, and the mechanosensitive *notch* (Smp_140800) (*77*), are consistent with the mechanical demands of a junction that transmits tail thrust to the body and severs as the cercaria penetrates the *stratum corneum* (the outer layer of mammalian skin).

**S1.4 Tail circular and longitudinal muscles** (2,122 cells)
*Low-density lipoprotein receptor* (*LDLR*, Smp_020550) was detected in 58% of cells and localised around ~100 nuclei along the lateral sides of the tail stem and forks, corresponding to the tail circular muscles, and four nuclei in the tail-stem midline, corresponding to the longitudinal tail-stem muscles (Fig. 1; fig. S4F). Signal at the body–tail junction reflects *LDLR* expression in 78% of junction-muscle cells.

**S1.5 Tail pseudostriated muscle** (363 cells)
*Acetyl-CoA hydrolase* (*ACH*, Smp_057060) was detected in 66% of cells and localised around ~57 nuclei in two columns flanking the tail longitudinal muscles, with no signal in the body or tail forks (Fig. 1; fig. S4F), matching previous descriptions based on ultrastructure and phalloidin staining (*27*, *50*, *78*). The cluster is enriched for the electron transport chain (BP), electron transfer activity (MF) and mitochondrial inner membrane (CC), consistent with the abundance of mitochondria described in these myocytes (*27*).

#### S2. Neurons

Six clusters expressed the neural markers *complexin* (*cpx*, Smp_050220), *prohormone convertase 2* (*PC2*, Smp_077980) and *Smp_068500*, a neural marker in adult worms (*11*). Segmentation of *cpx* ISH identified ~250 *cpx*+ nuclei, 195 in the body and 55 in the tail, so neurons account for 23% of body nuclei and 19% of tail nuclei (fig. S5A). Five clusters had markers specific enough for spatial validation.

**S2.1 *Prox-1*+ neurons** (111 cells)
*Homeobox protein prospero* (*prox-1*, Smp_045470) localised to two bilateral pairs of cells dorsal to the brain and beneath the body-wall muscles (Fig. 1; fig. S5C). Orthologues of several enriched genes function in eye specification and phototransduction: *meis* (Smp_174170), *otx* (Smp_124010), *six-related* (Smp_210120), *eya* (Smp_245330), *β-arrestin* (Smp_176980) and *r-opsin2* (Smp_104210; data S2) (*14*, *79*).

**S2.2 *SmKK7*+ neurons** (261 cells)
*UPF0506 domain-containing protein* (*UPF*, Smp_331460) and *SmKK7* (Smp_194830) were detected in nearly all cells. *UPF* localised to 10 nuclei: two in the distal tail stem and eight in the body, in three bilaterally symmetrical domains lateral to the acetabulum, dorsal to the brain and in the anterior organ (Fig. 1; fig. S5C), consistent with previous reports (*80*). The cluster is enriched for chemical synaptic transmission and G protein-coupled receptor signalling (BP), G protein-coupled receptor and voltage-gated potassium channel activity (MF) and presynapse (CC). The potassium-channel enrichment is notable because *SmKK7* encodes a proposed immunomodulator with homology to the potassium-channel blockers of scorpion venom (*81*). SAMap aligned the cluster with adult neuron 31 (7b2−), whose top markers are also *UPF* and *SmKK7* (fig. S2; data S2). *SmKK7* was first identified in the secretions of artificially transformed cercariae and assumed to be released by the acetabular glands, but its transcripts lie outside those glands and its protein localises to the peripheral nerve net and ciliated sensory endings (*80*). We detect *SmKK7* only in this cluster and the oesophageal gland (data S2), suggesting secretion across the tegument or into the oesophagus rather than from the acetabular glands.

**S2.3 *STK*+ neurons** (513 cells)
*Serine/threonine kinase* (*STK*, Smp_127060) localised to 22 bilaterally symmetrical cells in the body, at the periphery of the brain, near the head gland and beneath the dorsal and ventral body-wall muscles, and to ~11 nuclei along the tail midline (Fig. 1; fig. S5D). In both body and tail, labelled transcripts also lay away from *STK*+ nuclei, between and external to the body-wall muscles (fig. S5D). Other top markers, *G-protein α-i subunit* (*GNAI*, Smp_246100) and a *calmodulin* (Smp_343710), mark *GNAI*+ neurons in the schistosomulum (*12*), this cluster's reciprocal counterpart. SAMap also aligned it with a polyclad neuron in and around the brain (*MIP*+; fig. S2, data S2).

**S2.4 *Cornifelin*+ neurons** (669 cells)
*Cornifelin* (Smp_135250) was detected in 93% of cells and localised around ~42 nuclei in the tail midline, with weaker expression around ~8 nuclei in the body (Fig. 1; fig. S5D). The cluster is enriched for ciliary terms: cilium assembly, microtubule-based movement and intraciliary transport (BP), dynein light intermediate chain binding and minus-end-directed microtubule motor activity (MF), and cilium, axoneme and motile cilium (CC). SAMap aligned it with ciliated neurons in miracidia, adults and planarians (fig. S2, data S2), and immunolabelling with

acetylated tubulin showed cilia along the central length of the tail stem, which we hypothesise belong to these cells (fig. S5E). Other top markers support a ciliated sensory identity: a *TWIK-related potassium channel* (Smp_141570), *polycystin-1-related* (Smp_165230), whose *C. elegans* orthologue functions in contact sensing and localises to cilia, and the ciliated-neuron-enriched *P25-alpha-related* protein (Smp_097490).

**S2.5 *GPCR-B*+ neurons** (39 cells)
The smallest cluster in the atlas. *GPCR class B* (*GPCR-B*, Smp_099670), of the secretin/adhesion receptor family (*82*), localised to two cells posterior to the body–tail junction and one nucleus at the distal end of the tail stem (Fig. 1; fig. S5F). Two of its five top markers are neuropeptide precursors: *Smp_188580*, originally *SmNPP-23* (*83*) and later renamed *SmNPP-35* (*84*), and *Smp_204880*, found only in closely related species. Immunofluorescence with an antibody we generated against SmNPP-35, with phalloidin staining, placed these cells along the dorsal nerve tract of the tail beneath the longitudinal muscles (fig. S5F), suggesting a motor neuron identity. SAMap aligned the cluster with an undescribed adult neuron population (N17), peptidergic neurons in miracidia (Neuron 2) and planarian *pc2*+ neurons (fig. S2, data S2).

**S2.6 Undefined neural** (4,743 cells; not spatially validated)
No marker gene distinguished this cluster at atlas resolution, so we subclustered it, recovering 21 subclusters, 13 with distinct genes among their top three markers (fig. S6). Two had neuropeptide precursors as top markers: *neuropeptide Y* (*NPY*, Smp_159950), localised to two bilaterally symmetrical nuclei dorsal to the brain with projections running posteriorly between the longitudinal body-wall muscles to the body–tail junction, and *neuropeptide F* (*NPF*, Smp_088360), localised to a dorsal pair above the *NPY* cells, a pair below and lateral to the brain, and a pair lateral to the acetabulum (fig. S6). We infer that the remaining cells are also in the body, because every *cpx*+ nucleus counted in the tail was accounted for by the validated clusters above.

**S3. Tegument**

**S3.1 Body tegument** (1,335 cells)
Identified from tegument markers described in other life stages: *micro-exon gene 6* (*MEG-6*, Smp_163710), *tetraspanin-2* (*tsp-2*, Smp_335630), *glucose transporter-4* (*gtp-4*, Smp_105410) and *nucleotide pyrophosphatase/phosphodiesterase 5* (*npp-5*, Smp_153390) (*11*, *13*, *14*). We validated it with an uncharacterised gene, *Smp_057860*, detected in 92% of its cells and localised around ~33 nuclei in the body beneath the body-wall muscles, with none in the tail (Fig. 1; fig. S8A).

**S3.2 Anterior body tegument** (68 cells)
This cluster adjoins the body tegument but lacks the canonical markers *MEG-6*, *tsp-2* and *gtp-4*. We validated it with *DUF3421 domain-containing protein* (*DUF3421*, Smp_035550), detected in 43% of cells and localised to six cells within the muscular head cone, dorsal to the mouth and surrounding the head gland (Fig. 1; fig. S8A). Its top markers are relatively unspecific, including *calmodulin* (Smp_032970), *annexin* (Smp_074140) and a secreted protein (Smp_173610). Given the anterior position around the mouth and the absence of canonical tegument genes, we hypothesise that it is intermediate between tegument and digestive cell types and may be

associated with the head glands. It had the weakest SAMap alignments of any cercarial cell type to other life stages and flatworms, indicating a distinct transcriptome (fig. S2, data S2).

**S3.3 Tail tegument** (362 cells)
Marker gene *larval extracellular vesicle-1* (*LEV1*, Smp_074560) was detected in 76% of cells. This cluster is described in the main text.

**S4. Digestive**
In adult schistosomes the digestive tract comprises a mouth, an oesophagus whose anterior cell mass and oesophageal gland secrete factors that process ingested blood, and the syncytial gut (*85*). Although the cercaria does not feed, it has a non-functioning gut primordium (*86*).

**S4.1 Anterior oesophagus** (149 cells)
*Calciphosine-like protein* (*CAPS-like*, Smp_096390) localised around three nuclei posterior to the mouth, with transcripts surrounding the anterior oesophageal tract before it passes through the muscles of the head cone (Fig. 1; fig. S9, B and C). Other markers, *MEG-12* (Smp_152630) and *phospholipase A* (*PLA2*, Smp_031190), are expressed in the anterior oesophagus of the adult worm (*30*), alongside *MEG-28* (Smp_243760), *VAL-13* (Smp_124060) and *aquaporins 3* and *9* (Smp_300990, Smp_332980). Notably, *CAPS-like* is also the top marker of the miracidial ciliary plates (*14*), yet SAMap does not align this cluster with ciliary plates (fig. S2, data S2), highlighting that cell types should be compared transcriptome-wide rather than by marker genes alone.

**S4.2 Oesophageal gland** (238 cells)
*MEG-14* (Smp_124000) localised to 10 cells surrounding the posterior oesophagus (Fig. 1; fig. S9D). Eight further micro-exon genes were among the top markers (*MEG-4*, *-6*, *-8*, *-9*, *-11*, *-15*, *-18*, *-32*), as were two venom allergen-like genes (*VAL-7*, Smp_300070; *Smp_300080*) (fig. S9E, data S2). *MEG-14* binds calgranulin, a human S100 protein involved in inflammation (*87*). SAMap aligned the cluster with a planarian parenchymal population described as candidate gland cells (fig. S2, data S2).

**S4.3 Intestine** (67 cells)
*Lysozyme-associated membrane glycoprotein* (*LAMG*, Smp_162770) localised to around seven cells forming the intestine (Fig. 1; fig. S9D). Other top markers include *cathepsins B*, *D* and *L* (Smp_067060, Smp_103610, Smp_343260, Smp_085010, Smp_013040, Smp_105370) and saposin-containing proteins (Smp_105420, Smp_105450, Smp_194910, Smp_028870, Smp_181070) (fig. S9E, data S2). Cathepsins digest host haemoglobin, serum albumin and immunoglobulin G (*88–90*), and saposin-like proteins are candidate pore-forming haemolysins of the schistosome gut (*91*). Gut differentiation begins only in three-week-old juveniles (*92*), so this expression appears to anticipate the onset of blood feeding.

**S5. Excretory (protonephridial) and secretory cell types**
These are the first excretory tube cells of *S. mansoni* captured by scRNA-seq, and the body and tail tubes are unexpectedly distinct. We also recovered a previously undescribed tail secretory cell type.

**S5.1 Flame cells** (624 cells)
The cercaria has four pairs of flame cells in the body and one pair at the anterior of the tail, whose beating flagella draw in and filter interstitial fluid (*50*). The flame cell marker *ShKT-domain-containing protein* (*ShKT*, Smp_335600) (*93*) localised around the nuclei of all ten, identified by their proximity to the barrels of the tube cells, counterstained with phalloidin (Fig. 1; fig. S7, A and B). Its enriched terms are ciliary: cilium assembly, microtubule-based movement and intraciliary transport (BP), dynein light intermediate chain binding and minus-end-directed microtubule motor activity (MF), and cilium, axoneme and motile cilium (CC).

**S5.2 Body excretory tubes** (184 cells)
Tubule cells convey filtrate from the flame cells to a bladder at the posterior of the body (*86*). *Family M13 non-peptidase homologue* (*M13*, Smp_000755) localised to the bilaterally symmetrical tubule cells running between the four pairs of body flame cells, and extending into the tail to reach the pair of tail flame cells. *M13* is co-expressed with *clumping factor A* in the body tubes but is absent from the tail tubes (below) (Fig. 1; fig. S7, C and D).

**S5.3 Tail excretory tubules** (434 cells)
Two tubules pass down the centre of the tail, one entering each fork and terminating at an excretory pore (nephridiopore) at the tip (*86*). *Clumping factor A* (*CFA*, Smp_194050) was expressed in both tube clusters, localising to the body tubules, through the body–tail junction to the tail flame cells, and along the tubule cells of the tail stem and forks (fig. S7, D and E). Since no *M13+* nuclei are in the tail, the *CFA+* tail cells derive exclusively from this cluster, and co-labelling with an anti-phospho-tyrosine antibody, which marks the tail excretory duct (*50*), confirmed their identity (fig. S7E). Comparing the two excretory tube populations directly, the tail tubules are enriched for the electron transport chain (BP), electron transfer activity (MF) and mitochondrial inner membrane (CC), suggesting they need more energy to clear the local metabolic load of vigorous tail beating.

**S5.4 Secretory cells** (95 cells)
*Smp_331550* was detected in 93% of cells, in the lumen of the pre- and post-acetabular glands and in ~6 cells along the midline of the lower tail stem and fork (Fig. 1; fig. S7F). Co-staining with peanut agglutinin, which labels the acetabular glands (*50*), confirmed that the body signal corresponds to those glands (fig. S7G). Given their size (~23 × 103 µm) the acetabular glands are unlikely to have been captured in the 10x emulsion (maximum recommended cell diameter 40 µm), so the cluster transcriptome is attributed to the *Smp_331550+* tail cells. It expresses several genes encoding secreted proteins, including *MEG-7* (Smp_165050), *venom allergen-like 16* (*VAL-16*, Smp_124070) and uncharacterised secreted proteins (*Smp_323320*, *Smp_319290*, *Smp_323440*, *Smp_323230*). SAMap aligned it with planarian parenchyma subcluster 10 (fig. S2, data S2), which contains the marginal adhesive gland cells (*15*).

**S6. Parenchyma (772 cells)**
*Serpin* (Smp_090080) was detected in 98% of cells and localised to 24 cells in the body (Fig. 1; fig. S10A), which have long cytoplasmic protrusions reaching between surrounding cells and filling the intercellular space. The cluster is enriched for *glycogen branching enzyme* (Smp_094880) and *glycogen phosphorylase* (Smp_143840), consistent with the glycogen granules described in these cells (*86*). Their restriction to the body implies that tail cells draw on

another store, most likely the pseudostriated muscle, which also holds large amounts of glycogen (*86*).

**S7. Stem cells (366 cells)**

Identified from stem-cell markers of other life stages: *ago2-1* (Smp_179320), *nanos2* (Smp_051920), *p53* (Smp_139530), *fgfrA* (Smp_175590), *fgfrB* (Smp_157300) and *hesl* (Smp_024860); fig. S10B) (*14*, *94*, *95*). *Linker histone H1* (*LH1*, Smp_162370), detected in 83% of cells, localised around ~12 nuclei in the body: two in the dorsal midline, two bilaterally symmetrical pairs on the lateral sides, four around the acetabulum and two at the posterior (Fig. 1; fig. S10, C and D). *ago2-1* and *nanos2* gave the same body pattern; *ago2-1* was additionally expressed at low level in flame cells and tail excretory tubules in silico, and in two to three further cells of the tail stem *in situ* (fig. S10E). Although stem cells only begin dividing one day after transformation (*92*), the cercarial cluster is already enriched for DNA replication and DNA replication initiation (BP), structural constituent of chromatin (MF) and nuclear chromosome (CC). Unlike the miracidium, which has two stem populations further separated by sex (*14*, *94*, *95*), the cercaria has a single stem cluster containing both sexes (fig. S1C).

**S8. Ambiguous (2,812 cells)**

This cluster lacked marker genes specific enough for spatial validation and did not resemble any known cell type or tissue, showing instead non-specific expression of muscle and neural markers throughout. It may represent cells in transition, or lower-quality cells that were damaged, stressed or affected by technical factors such as gene dropout; we did not attempt ISH validation given the absence of robust markers. Expression of the putative stem-associated genes *Smp_034860* and *Smp_323010* in a subset of cells suggests it may also contain early or precursor stem-like cells.

### Supplementary Figures

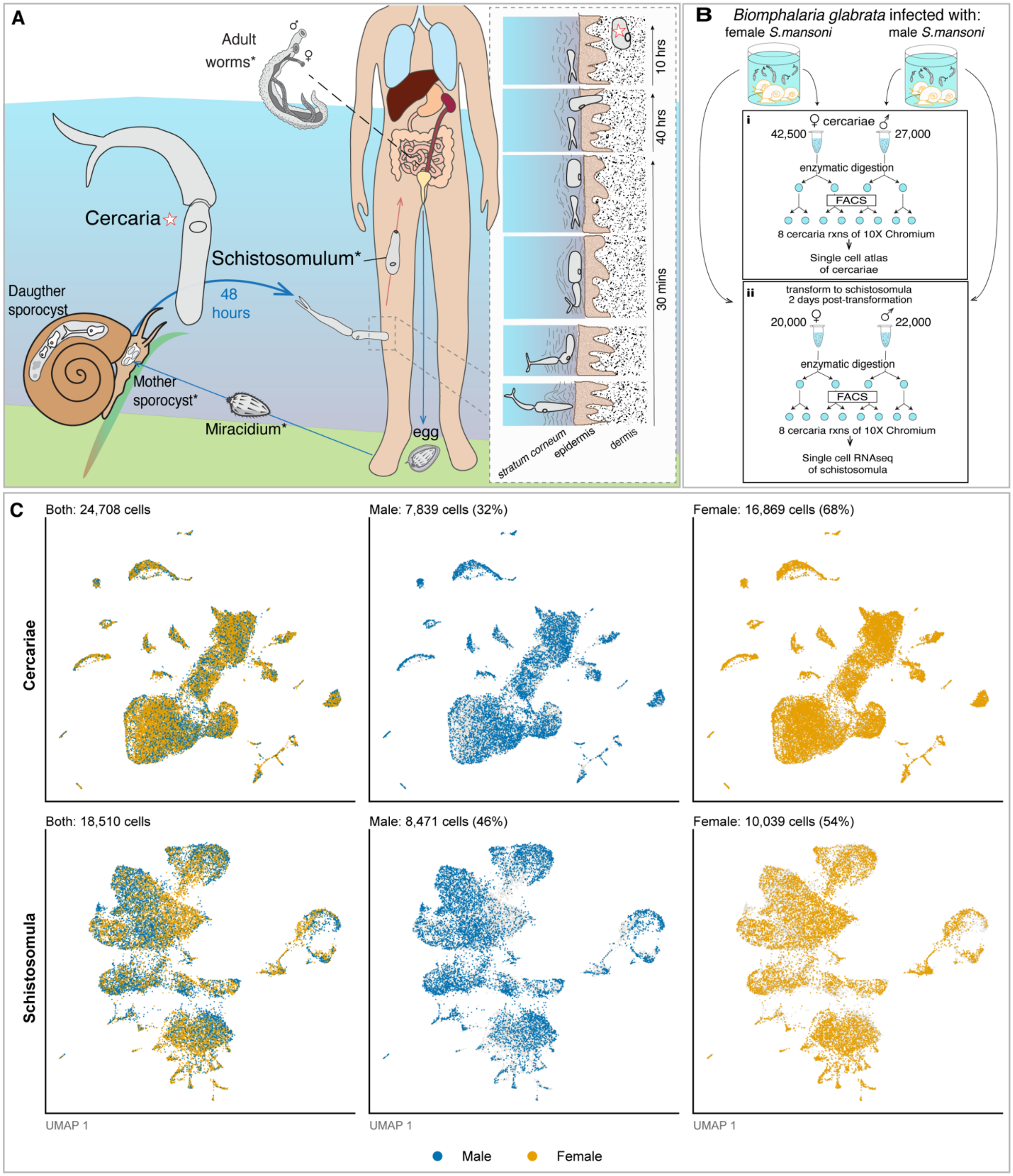


**Fig. S1. Overview of *Schistosoma mansoni* life-cycle stages collected for single-cell RNA sequencing (scRNA-seq). (A)** Schematic of the complex life cycle of *S. mansoni*, including a timeline of cercarial transmission and infection (*4*, *5*, *96*). Asterisks, life-cycle stages with published whole-body single-cell atlases; ☆, scRNA-seq data generated in this study. **(B)** Experimental scheme showing the *Biomphalaria glabrata* intermediate host snail infected with either male or female *S. mansoni*. Cercariae were shed from snails, and male and female cercariae were kept separate for (i) dissociation into single-cell suspensions for scRNA-seq to generate a cercaria atlas and (ii) mechanical transformation into schistosomula and two days of *in vitro* culture before dissociation into single-cell suspensions for scRNA-seq to generate a schistosomulum atlas. **(C)** UMAPs of cercarial and schistosomulum clusters showing male and female cells.

| *S. mansoni* Cercaria | | *S. mansoni* Life Stages | | | Other Flatworms | | Mean Alignments | | |
|---|---|---|---|---|---|---|---|---|---|
| Category | Cluster | Miracidia | Schistosomula | Adult | *S. mediterranea* | *P. crozieri* | *S. mansoni* | Other Flatworms | All |
| Muscle | Body circular & acetabular anchor | Stem G (0.52) | *myoD+* muscle (0.79) | Muscle 5 (0.24) | Parenchyma: 11 (0.24) | Neo-1 (0.24) | 0.51 | 0.24 | 0.40 |
| | Body longitudinal | Muscle 1 (0.46) | pou6f1/brn-5+ muscle (0.33) | Muscle 2 (0.85) | Muscle: 6 (0.56) | Myo-1 (0.30) | 0.55 | 0.43 | 0.50 |
| | Body-tail junction | Muscle 1 (0.41) | Positional muscle (0.88) | Muscle 3 (0.02) | Muscle: 4 (0.08) | Myo-1 (0.18) | 0.44 | 0.13 | 0.31 |
| | Tail circular & longitudinal | Muscle 1 (0.66) | Positional muscle (0.36) | Muscle 4 (0.63) | Muscle: 12 (0.62) | Myo-2 (0.77) | 0.55 | 0.69 | 0.61 |
| | Tail pseudostriated | Muscle 2 (0.23) | *myoD+* muscle (0.08) | Muscle 1 (0.08) | Muscle: 9 (0.10) | Gut-1 (0.15) | 0.13 | 0.12 | 0.13 |
| Neural | *Prox-1+* | Neuron 1 (0.07) | *7b2/pc2+* neurons (0.20) | Neuron 10 (0.03) | *opsin+* Neural (0.50) | Neuron-*GNRH+* (0.16) | 0.10 | 0.33 | 0.19 |
| | *SmKK7+* | Neuron 1 (0.13) | *SmKK7+* neurons (0.85) | Neuron 31 (7b2-) (0.89) | Parenchyma: 11 (0.38) | Neuron-*Sy65-4+* (0.15) | 0.63 | 0.27 | 0.48 |
| | *STK+* | Neuron 5 (0.36) | *gnai+* neurons (0.89) | Neuron 7 (0.19) | Neural: 10 (0.35) | Neuron-*MIP+* (0.49) | 0.48 | 0.42 | 0.46 |
| | *Cornifelin+* ciliated | Neuron 4 (0.83) | *7b2/pc2+* neurons (0.44) | Neuron 30 (0.54) | Neural: Ciliated: 5 (0.71) | Cilia-3 (0.22) | 0.60 | 0.47 | 0.55 |
| | *GPCR+* | Neuron 2 (0.78) | *7b2/pc2+* neurons (0.56) | Neuron 17 (0.85) | Neural: 3 (0.50) | *Neuron-GNRH+* (0.16) | 0.73 | 0.33 | 0.57 |
| | Undefined Neural | Neuron 1 (0.83) | *eag+* neurons (0.96) | Neuron 18 (0.89) | Neural: 3 (0.81) | Neuron-cilia-2 (0.54) | 0.89 | 0.68 | 0.81 |
| Tegument | Body tegument | Tegument (0.69) | Tegument (0.75) | Tegument 1 (0.85) | Pharynx: 6 (0.86) | Cilia-1 (0.66) | 0.76 | 0.76 | 0.76 |
| | Anterior body tegument | Parenchyma 2 (0.06) | Excretory tubes (0.07) | Tegument 1 (0.17) | Epidermal: 5 (0.04) | Gut-1 (0.07) | 0.10 | 0.06 | 0.08 |
| | Tail tegument | Ciliary plate (0.04) | Non-specific muscle (0.17) | GSC progeny (0.30) | Epidermal: 0 (0.07) | *Mes-TegA+*1 (0.43) | 0.17 | 0.25 | 0.20 |
| Digestive | Anterior oesophagus | Tegument (0.18) | *actin-2+* muscle (0.06) | *Meg-1+* (0.05) | Epidermal: 0 (0.61) | Rha-2 (0.25) | 0.10 | 0.43 | 0.23 |
| | Oesophageal gland | Stem F (0.02) | Oesophageal (0.95) | Sm13+ (0.18) | Parenchyma: 7 (0.84) | 96 (0.47) | 0.38 | 0.66 | 0.49 |
| | Intestine | Stem F (0.07) | Intestine (0.99) | Gut (0.28) | Intestine: 4 (0.69) | 100 (0.27) | 0.45 | 0.48 | 0.46 |
| Excretory | Flame cells | Protonephridia (0.92) | Flame cells (0.95) | Flame cells (0.87) | Flame Cell: 1 (0.74) | Cilia-2 (0.26) | 0.91 | 0.50 | 0.75 |
| | Body excretory tubes | Parenchyma 2 (0.58) | Excretory tubes (0.97) | Tegument 2 (0.27) | Epidermal: 5 (0.26) | Proneph (0.12) | 0.61 | 0.19 | 0.44 |
| | Tail excretory tubes | Parenchyma 2 (0.82) | Excretory tubes (0.34) | Tegument 2 (0.09) | Epidermal: 11 (0.25) | *Mes-Macif+*1 (0.24) | 0.42 | 0.24 | 0.35 |
| | Secretory cells | Ciliary plate (0.16) | Non-specific muscle (0.09) | Mehlis' gland (0.34) | Parenchyma: 10 (0.94) | Rha-2 (0.50) | 0.20 | 0.72 | 0.41 |
| Support | Parenchyma | Parenchyma 1 (0.90) | Parenchyma (0.91) | Parenchyma 1 (0.90) | *Cathepsin+* cells: 5 (0.67) | *Cath+*-2 (0.62) | 0.90 | 0.65 | 0.80 |
| | Stem | Stem D (0.48) | Stem (0.94) | Neoblast 1 (0.51) | Neoblast: Neural 2: 5 (0.35) | Neo-1 (0.85) | 0.64 | 0.60 | 0.63 |
| | Ambiguous | Stem F (0.39) | Stem-like (0.78) | *Hes2+* (0.43) | Parenchyma: 2 (0.39) | Neuron-p1 (0.39) | 0.53 | 0.39 | 0.48 |

**Fig. S2. Highest cell-type alignment scores between the cercaria and other life-cycle stages and flatworms.** Each cercarial cell type was aligned with SAMap to the cell types of the miracidium (*14*), the 2-day schistosomulum (this study) and the adult (*11*), and of the free-living flatworms *Schmidtea mediterranea* (*15*) and *Prostheceraeus crozieri* (*16*). The life stages were aligned in one joint analysis and the flatworms in a second. Color, alignment score; text, the top-aligning cell type with its score in parentheses. All alignment scores are in data S2.

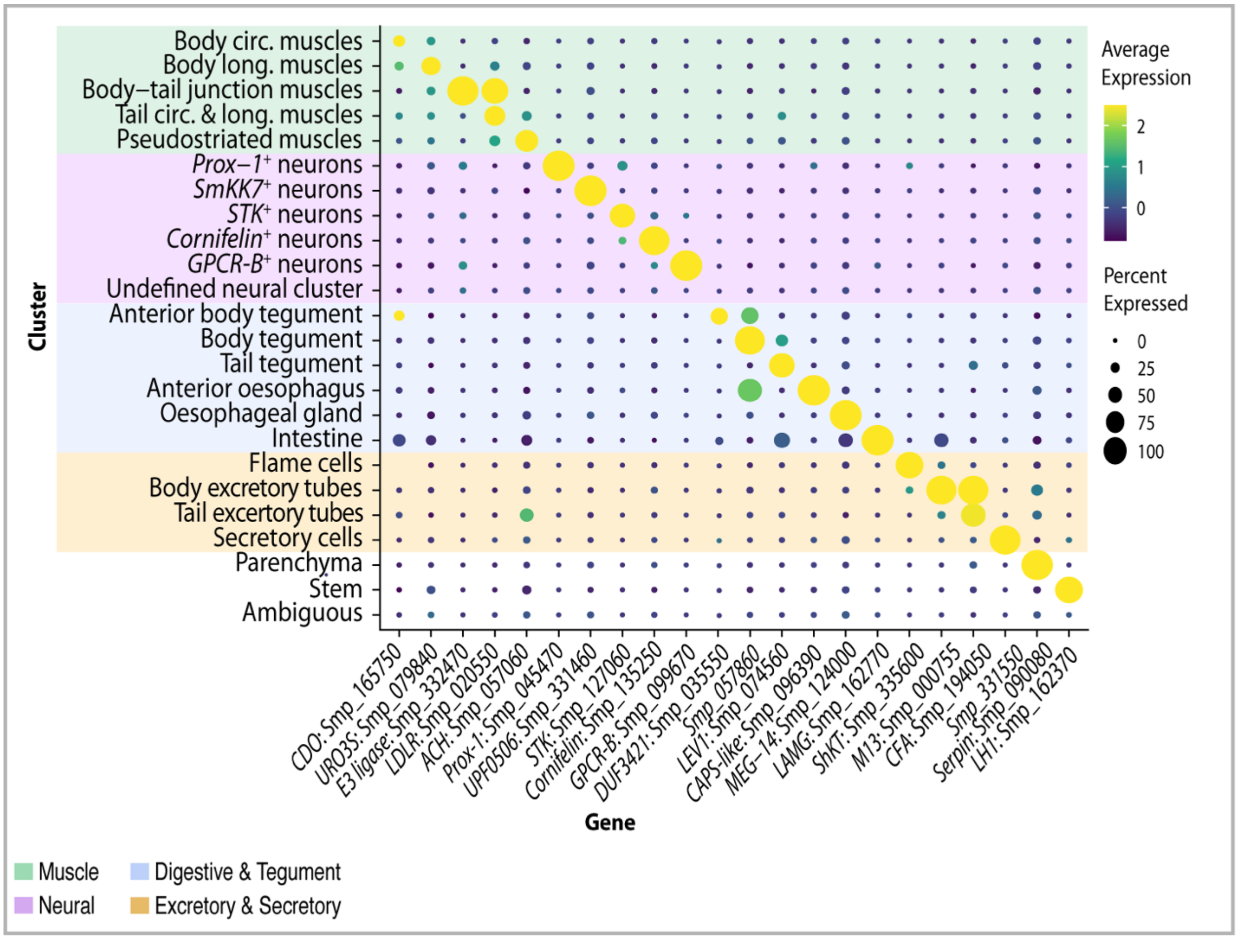


**Fig. S3. Identification of cluster marker genes for the cercarial cell types.** Dot plot of selected marker genes validated by *in situ* hybridization (ISH) for each cell type cluster. Color, average expression (yellow, highest; blue, lowest); dot size, percentage of cells in the cluster expressing the gene.

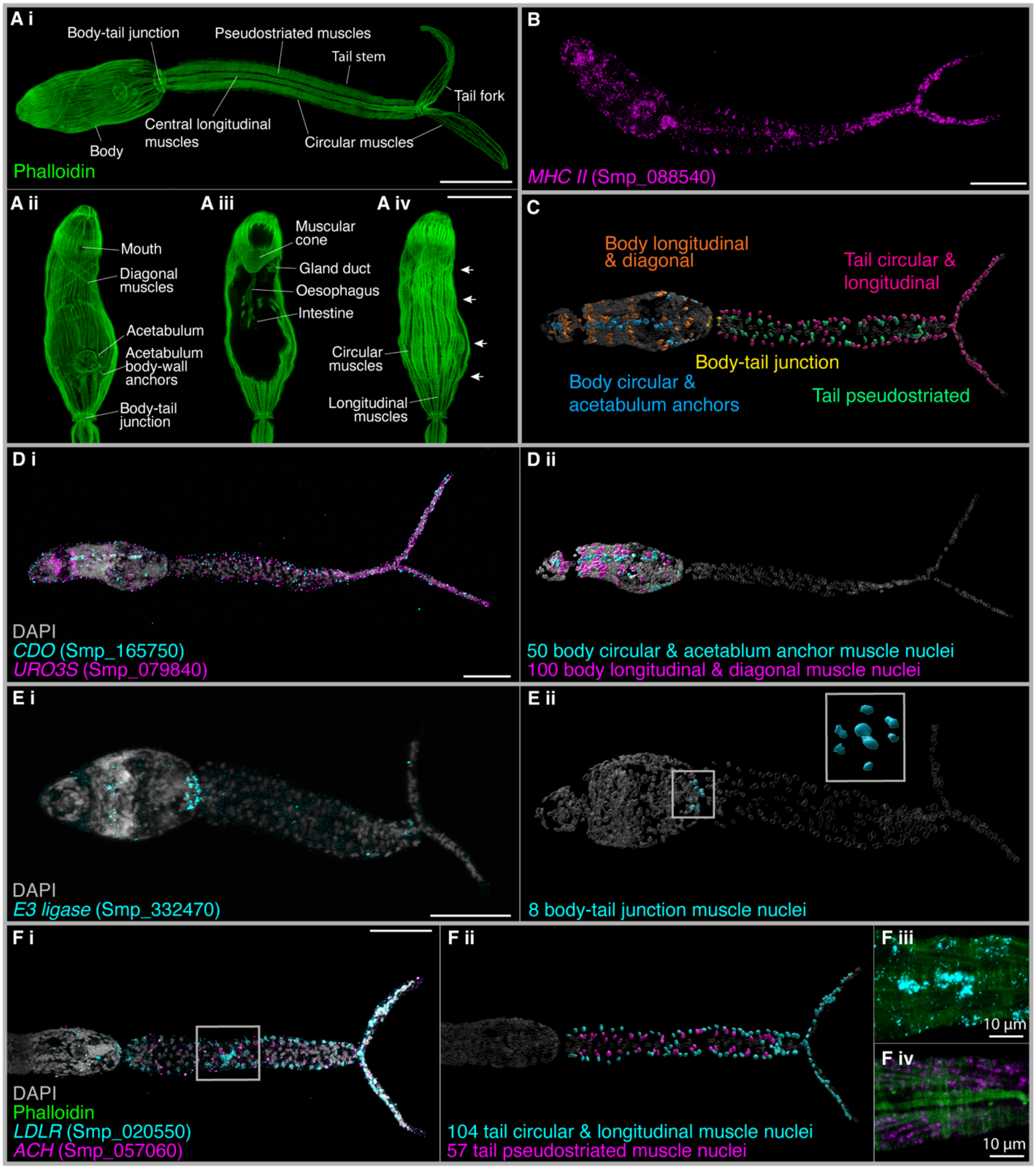


**Fig. S4. Transcriptionally distinct body and tail muscles. (Ai)** Maximum projection of phalloidin-labeled F-actin highlights the cercarial musculature. Ventral (Aii), midsection (Aiii) and dorsal (Aiv) planes of the body. Arrows, interdigitating tiers of longitudinal muscle. **(B)** Hybridization chain reaction (HCR) of muscle marker *MHC II*. **(C)** 3D model of muscle nuclei from two overlaid individuals (IMARIS segmentation). **(Di)** HCR of *CDO* and *URO3S* labels body wall muscle. **(Dii)** Segmented body wall muscle nuclei: ~50 (45–55) along the dorsal–ventral midline form the circular muscles, and ~100 (96–107) align with the five tiers of longitudinal muscle. **(Ei)** HCR of *E3 ligase* labels body–tail junction muscles. **(Eii)** Eight segmented *E3 ligase*+ nuclei; inset, the ring of six nuclei in the body and two central tail nuclei. **(Fi)** HCR of *LDLR* and *ACH* labels tail muscle nuclei. **(Fii)** ~104 nuclei (98–110) at the sides and center of the tail form the circular and longitudinal muscles, and ~57 (56–58) in a column form the pseudostriated muscles. **(Fiii and Fiv)** Boxed region in Fi with phalloidin: alternating circular-muscle nuclei around the tail stem (Fiii) and diagonal pseudostriated muscles (Fiv). Nuclei counts, range in five cercariae. Scale bars, 50 μm unless stated.

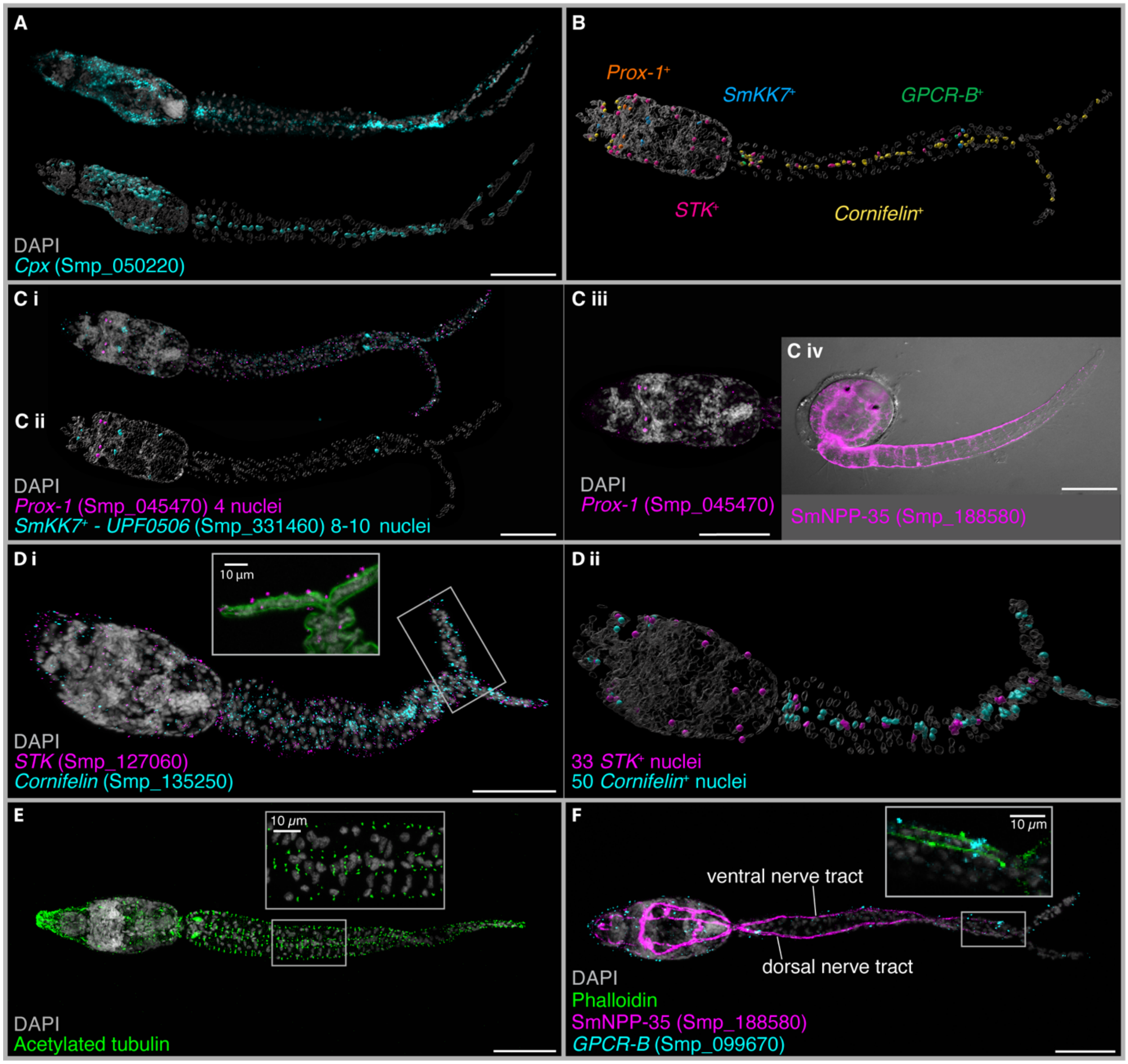


**Fig. S5. Cercarial neural diversity. (A)** HCR and segmentation of pan-neural marker *complexin* (*cpx*): 195 (190–203) positive nuclei in the body and 55 (54–55; three cercariae) in the tail. **(B)** 3D model of segmented nuclei expressing markers of the validated neural clusters. **(Ci)** HCR of *prox-1* and *UPF0506* shows body-specific *prox-1*+ neurons, and *SmKK7*+ neurons in the body and tail. **(Cii)** Segmented nuclei: four dorsal *prox-1*+ and 10 *SmKK7*+. **(Ciii)** *prox-1*+ cells lie dorsal to the brain, where the pigmented eyespots sit in the cercaria of *Lepocreadium setiferoides*. **(Civ)** DIC image overlaid with anti-SmNPP-35 immunofluorescence labeling the nervous system. **(Di)** HCR of *STK* and *cornifelin*. **(Dii)** Segmented nuclei: *STK*+, 22 (18–26) in the body and 11 (8–14) in the tail; *cornifelin*+, 8 (6–10) in the body and 42 (40–48) in the central tail stem. **(E)** Acetylated tubulin immunolabeling shows cilia enriched in the center and lateral sides of the tail stem and forks. **(F)** HCR of *GPCR-B* with SmNPP-35 immunolabeling: three *GPCR-B*+ nuclei align with the central dorsal nerve tract of the tail, between the central longitudinal tail stem muscles. Nuclei counts, range in five cercariae unless stated. Scale bars, 50 µm unless stated.

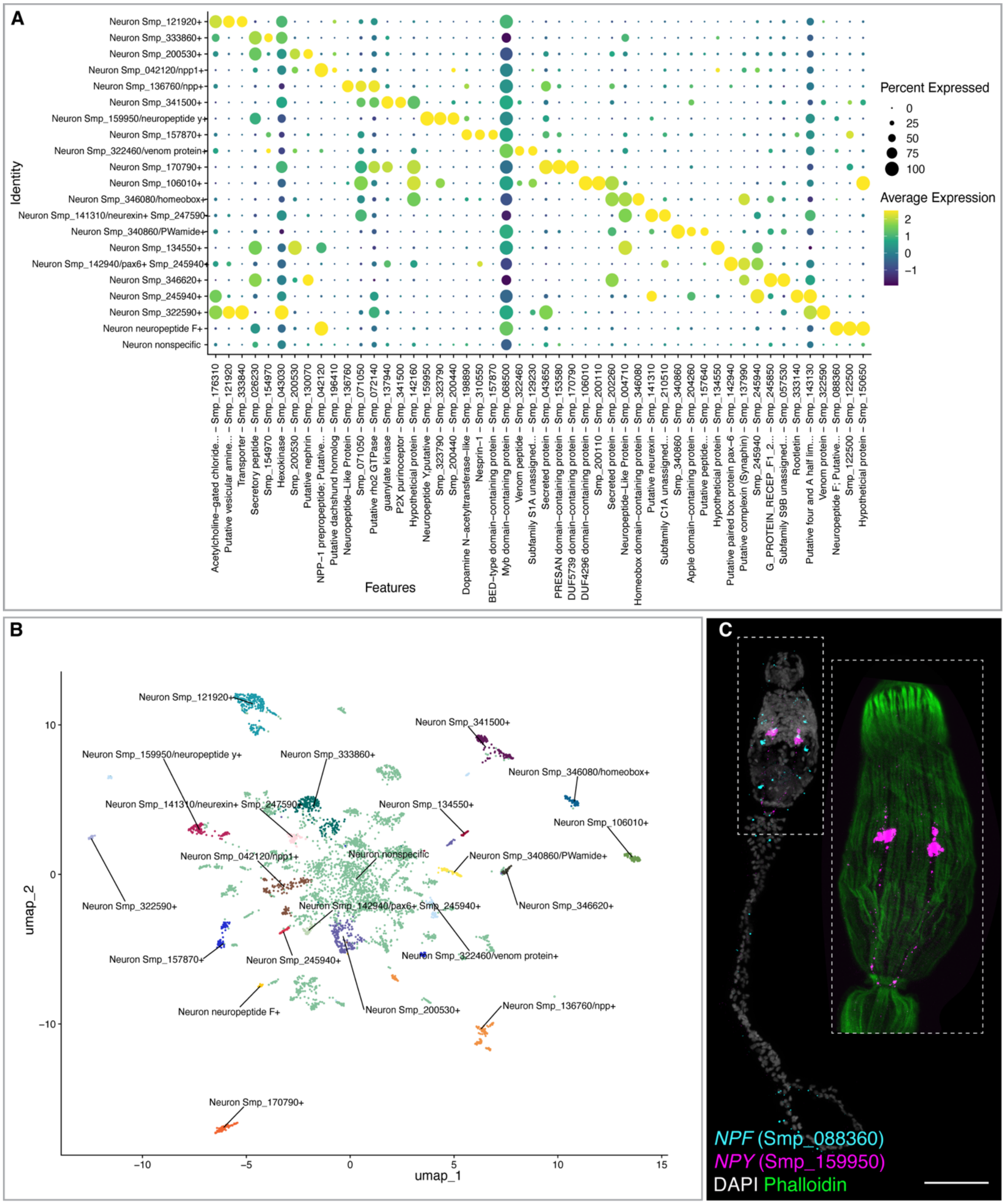


**Fig. S6. Undefined neural cluster includes a diversity of subpopulations. (A)** The undefined neural cluster was further divided into 21 subclusters; 13 have strong marker genes, as shown by their distinct expression in this dot plot of the top three marker genes of each subcluster. **(B)** UMAP of subclusters shows that many are distinct, while some remain adjoined. **(C)** Multiplexed HCR of *neuropeptide F* (*NPF*) and *neuropeptide Y* (*NPY*), top markers of two subclusters, localizes to the brain area. *NPY* expression also extends to the body–tail junction, running parallel to the longitudinal body wall muscles. Scale bar, 50 µm.

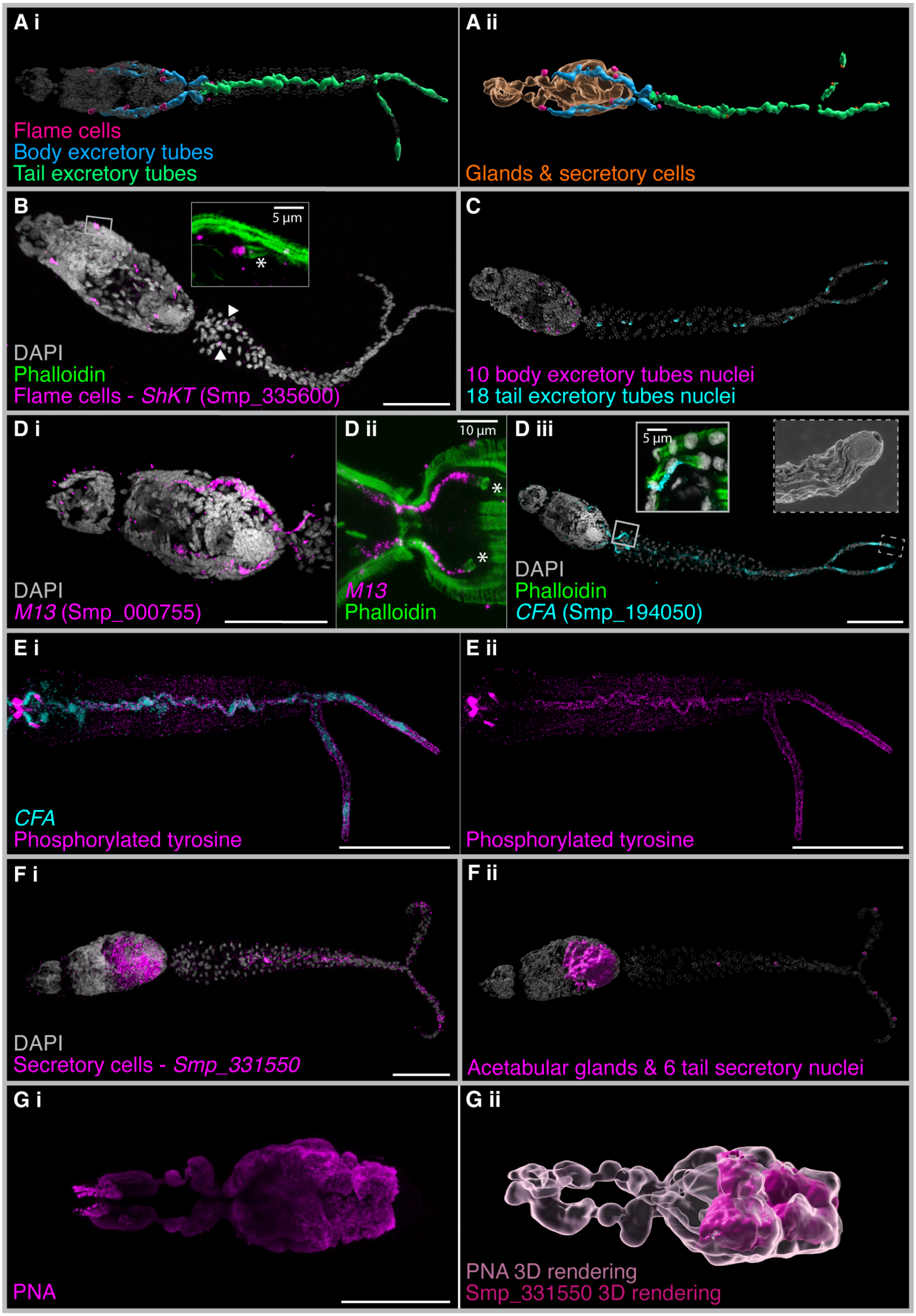
A i
Flame cells
Body excretory tubes
Tail excretory tubes
A ii
Glands & secretory cells
B
5 µm
DAPI
Phalloidin
Flame cells - ShKT (Smp_335600)
C
10 body excretory tubes nuclei
18 tail excretory tubes nuclei
D i
DAPI
M13 (Smp_000755)
D ii
10 µm
M13
Phalloidin
D iii
5 µm
DAPI
Phalloidin
CFA (Smp_194050)
E i
CFA
Phosphorylated tyrosine
E ii
Phosphorylated tyrosine
F i
DAPI
Secretory cells - Smp_331550
F ii
Acetabular glands & 6 tail secretory nuclei
G i
PNA
G ii
PNA 3D rendering
Smp_331550 3D rendering

**Fig. S7. Cercarial excretory and secretory cell types. (Ai)** Excretory system 3D model, ventral view: body tubes (blue) extend from flame cells (pink); tail tubes (green) run to the tail-fork tips. **(Aii)** Dorsolateral view: acetabular glands and secretory tail cells (orange). **(B)** HCR of flame cell marker *ShKT* in all 10 flame cells (weak in tail flame cells, ▼); inset, phalloidin labels the flame cell barrel (*). **(C)** Segmented tube nuclei: 10 body (magenta) and 18 tail (cyan). **(Di)** HCR of body tube marker *M13*. **(Dii)** *M13*+ cells extend through the body–tail junction to the tail flame cells (*). **(Diii)** Pan-tube marker *CFA* is coexpressed with *M13* in the body tubes (inset, solid line); *CFA*+ *M13*– tail cells are the tail tube cells, reaching the nephridiopores (inset, dashed line). **(E)** HCR of *CFA* with anti-phosphotyrosine immunolabeling confirms that tail *CFA*+ cells are tail tubes. **(Fi)** Secretory cell marker *Smp_331550* in six tail cells and the acetabular gland lumen. **(Fii)** 3D rendering of *Smp_331550*+ tail nuclei and gland-lumen signal. **(Gi)** Peanut agglutinin (PNA) labels the pre- and postacetabular glands. **(Gii)** Gland rendering (light pink) overlaid with *Smp_331550* (dark pink): colocalization in gland lumens. Scale bars, 50 µm unless stated.

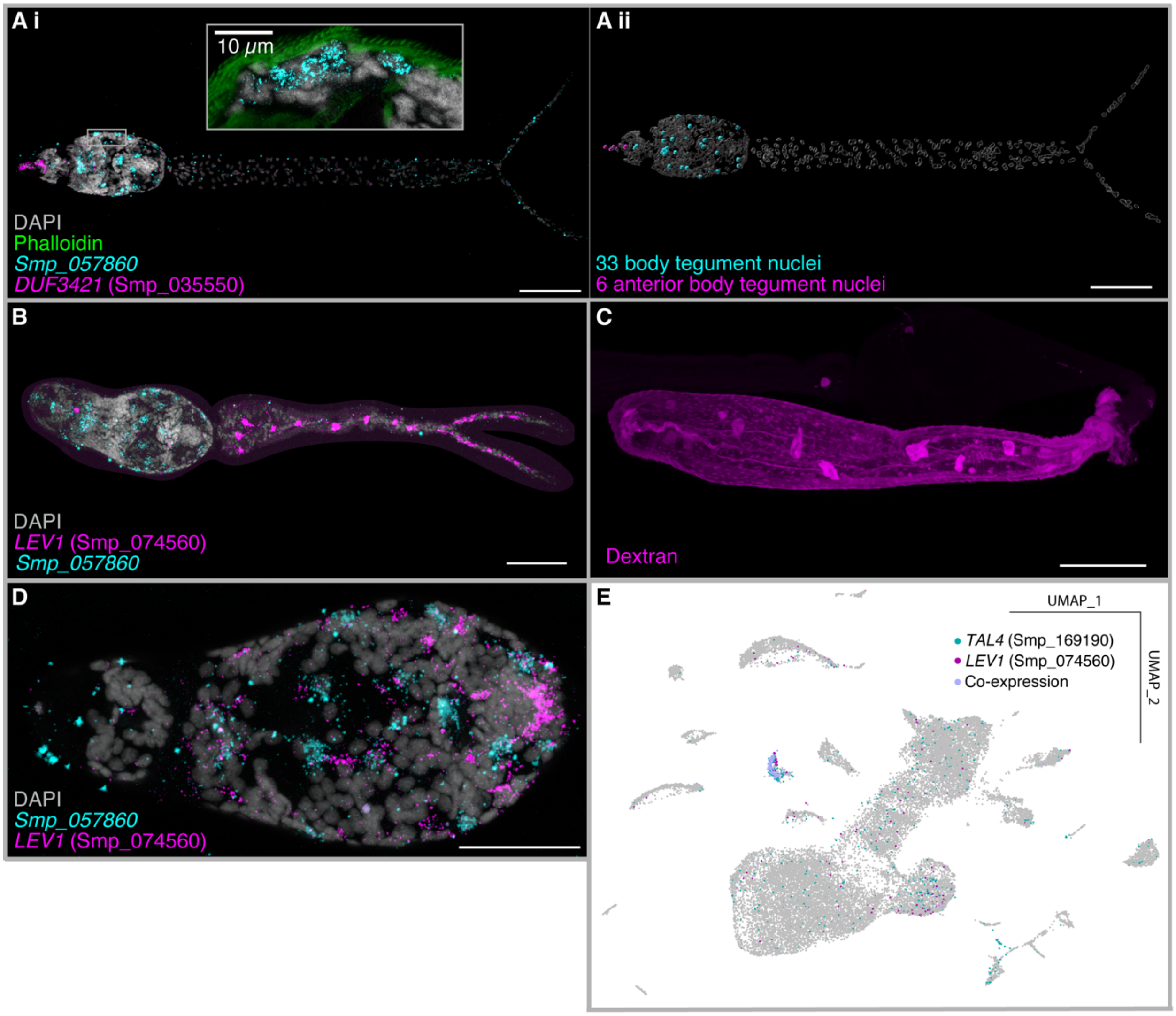


**Fig. S8. A previously undescribed, transcriptionally and morphologically distinct tail tegument. (Ai)** Multiplexed HCR of body tegument marker *Smp_057860* and anterior body tegument marker *DUF3421* localizes body tegument cells. Inset, phalloidin counterstaining reveals *Smp_057860* expression beneath the body wall muscles. **(Aii)** 3D rendering of segmented marker gene+ nuclei highlights 39 body tegument nuclei in total (32–34, *n* = 5; 5–6, *n* = 5). **(B)** Tail tegument marker *LEV1* is expressed in large domains down the tail stem and fork. **(C)** Dextran labeling shows patterns similar to *LEV1* expression in the tail stem. **(D)** *LEV1* is weakly expressed in the body, close to expression of the body tegument marker *Smp_057860*. **(E)** *LEV1* is highly coexpressed with *TAL4*, a known tail-specific gene (*19*). Scale bars, 50 µm.

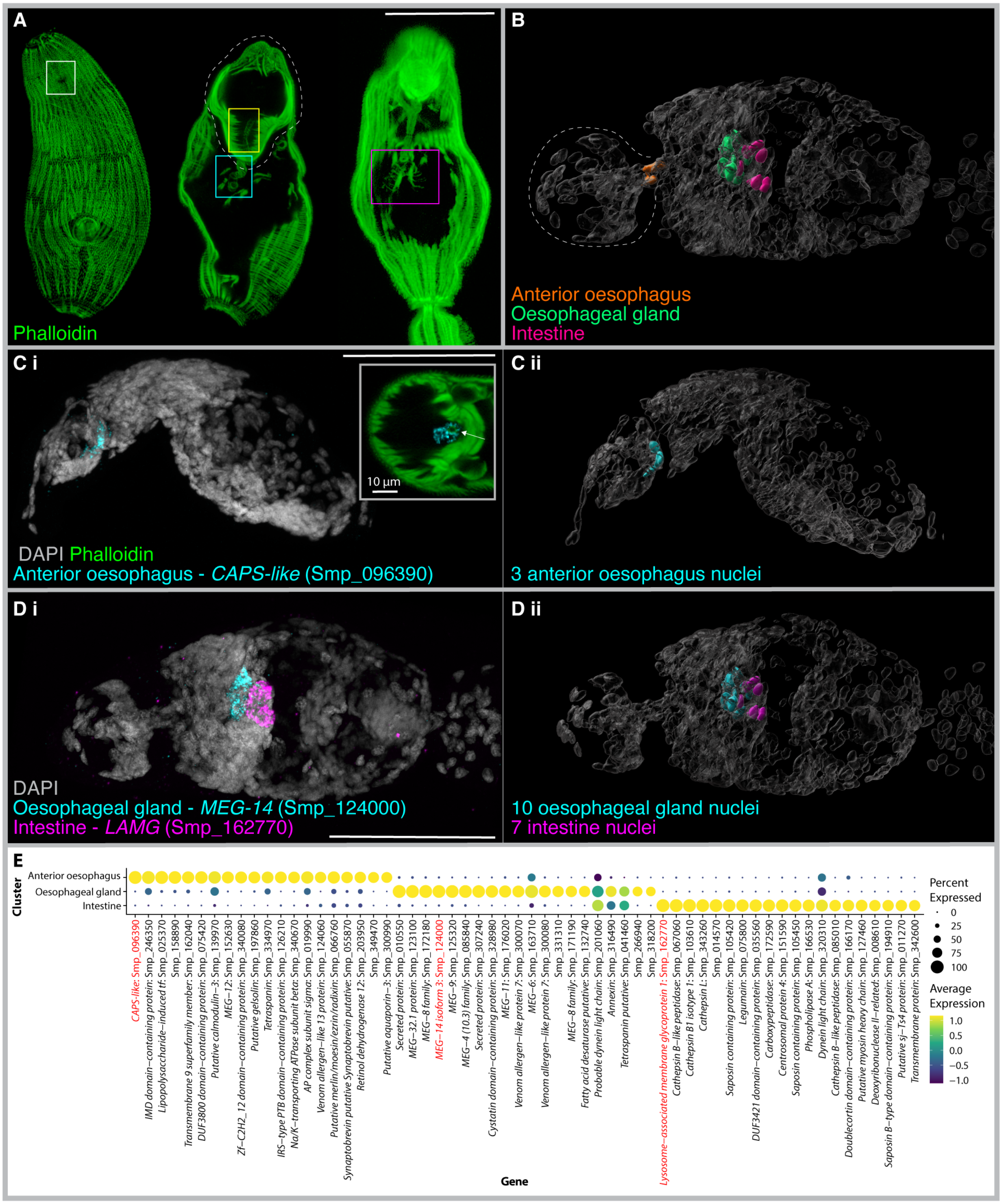


**Fig. S9. Cell types of the pre-functioning digestive system.** **(A)** Phalloidin labeling reveals the anatomy of the digestive system, including the mouth (white box), anterior oesophagus (yellow box) in the anterior organ (dashed line), oesophageal gland (blue box) and bifurcating intestine (magenta box). **(B)** 3D model of the digestive nuclei shows the anterior oesophagus (orange) at the base of the anterior organ (dashed line) and the oesophageal gland (green) anterior and adjacent to the intestine (magenta). **(Ci)** HCR of the anterior oesophagus marker gene *CAPS-like*. Inset, cross-section with phalloidin shows transcripts surrounding the anterior oesophagus (arrow). **(Cii)** 3D rendering of *CAPS-like*+ nuclei highlights three anterior oesophagus nuclei ($n$ = 5) on the ventral side of the anterior organ. **(Di)** Multiplexed HCR of oesophageal gland marker *MEG-14* and intestine marker *lysozyme-associated membrane glycoprotein*. **(Dii)** 3D rendering shows 10 oesophageal gland nuclei (*n* = 5) and seven intestine nuclei (*n* = 5) on the dorsal side, mid-body. **(E)** Top marker genes of the three digestive clusters include multiple MEG, VAL and secreted protein genes. Scale bars, 50 µm unless stated.

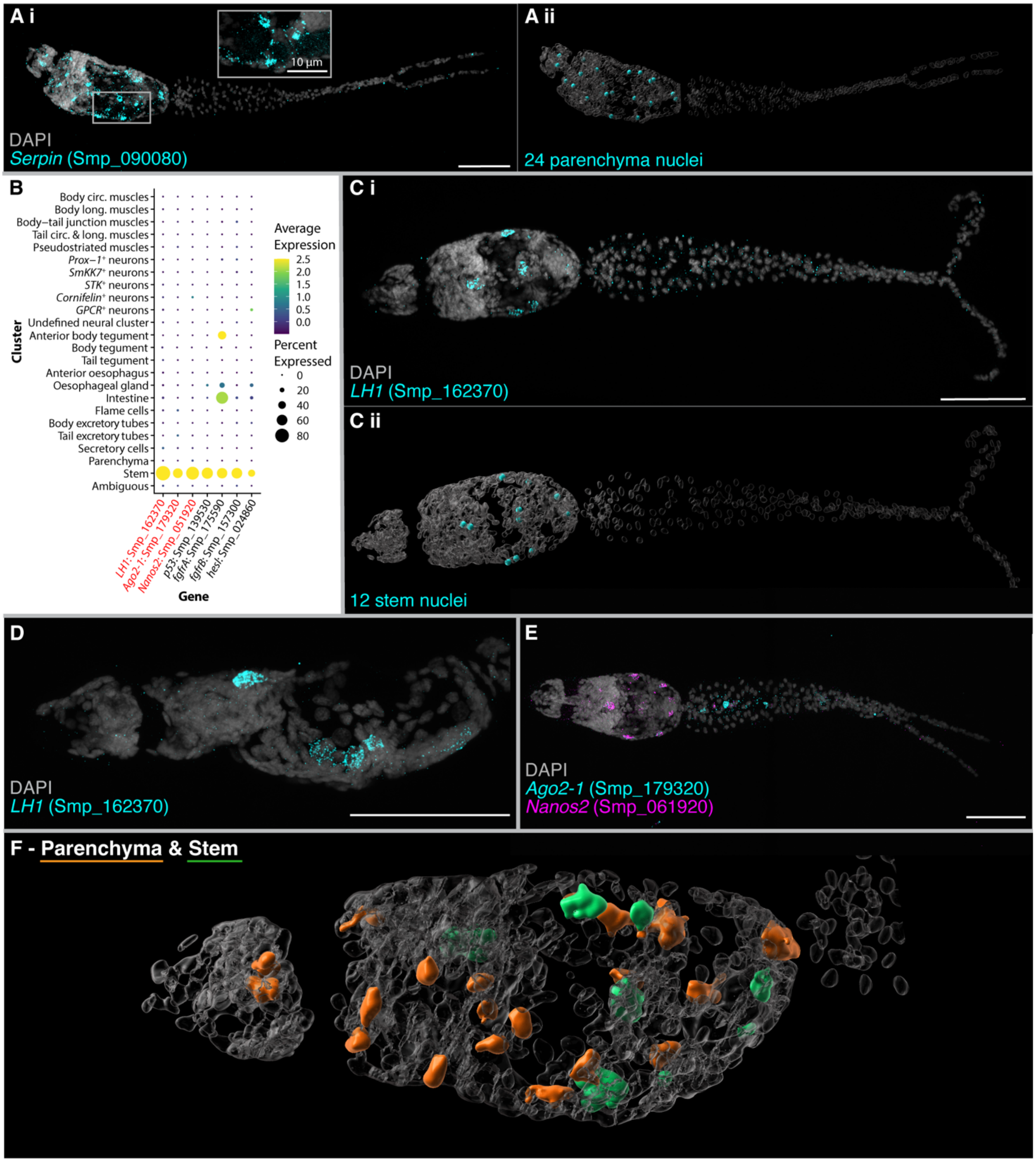


**Fig. S10. Parenchyma and stem cells are located in the body. (Ai)** HCR of parenchyma marker *serpin* shows expression in the body only. These *serpin*+ cells have cytoplasmic protrusions that extend to surrounding cells (inset). **(Aii)** 3D rendering of nuclei shows the spatial distribution of the parenchyma cells ($n = 3$). **(B)** Dot plot showing enrichment of known stem cell genes in the stem cluster. Genes validated by HCR are in red. **(Ci)** HCR of the stem cluster marker *LH1* localizes stem cells to the body only. **(Cii)** 3D rendering of segmented nuclei highlights 12 stem nuclei in the body. **(D)** Lateral view of *LH1* shows the dorsal and ventral localization of stem cells. **(E)** Multiplexed HCR of known stem markers *ago2* and *nanos2* shows expression patterns similar to *LH1* in the body, as well as *ago2*+ cells in the anterior tail stem. **(F)** 3D rendering of segmented marker gene expression of stem and parenchyma cells shows relative cell type positions. Scale bars, 50 µm unless stated.

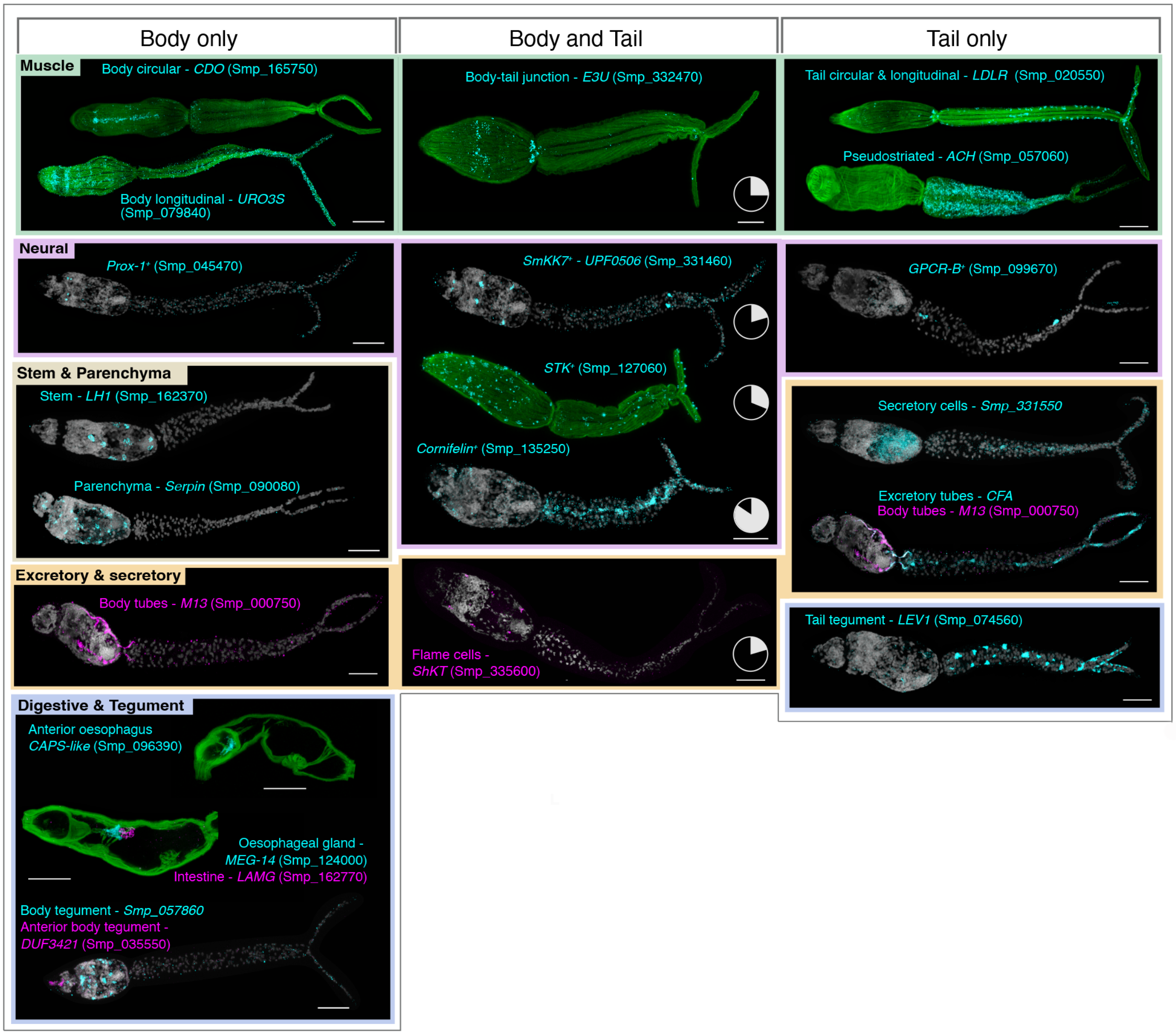


**Fig. S11. Differential distribution of cell types between the cercarial body and tail.** Within the cercaria, 22 clusters of cells were defined and spatially located. Of these, 11 are specific to the body, five span the body and tail, and six are tail specific. For cell types found in both (middle column), the white slice in the pie chart indicates the proportion of marker-expressing cells localized to the tail. The body excretory tubes (*M13*+, left column) extend into the tail, but their nuclei are in the body (fig. S7). In the right column, the marker for tail excretory tubes (*CFA*) is also expressed in the body tubes; *in situ* coexpression with the body tube marker shows that the *CFA*+/*M13*– cells are the tail tubes.

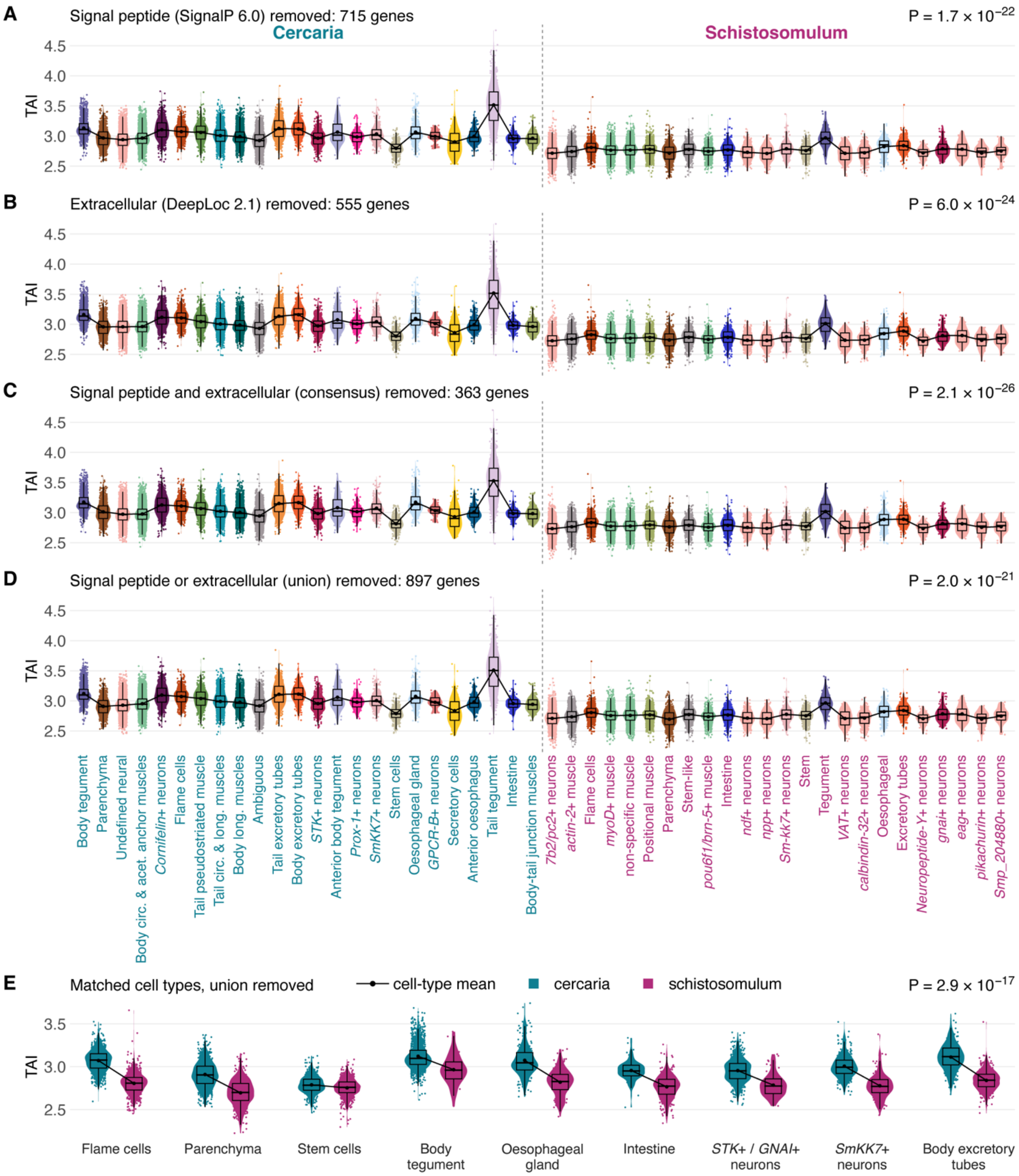


**Fig. S12. Cell-type TAI profiles after removing genes for predicted secreted or surface proteins.** Transcriptomic age index (TAI) of each cercarial and 2-day schistosomulum cell, recomputed without genes predicted to encode secreted or surface proteins: **(A)** signal peptide (SignalP 6.0; 715 genes), **(B)** extracellular (DeepLoc 2.1; 555 genes), **(C)** both (363 genes) and **(D)** either (897 genes). Gene numbers count only genes present in the atlas data; full lists in data S3. **(E)** The nine matched cercaria–schistosomulum cell-type pairs with the union set (D) removed, colored by stage. *P* values, myTAI flat line test across cell types.

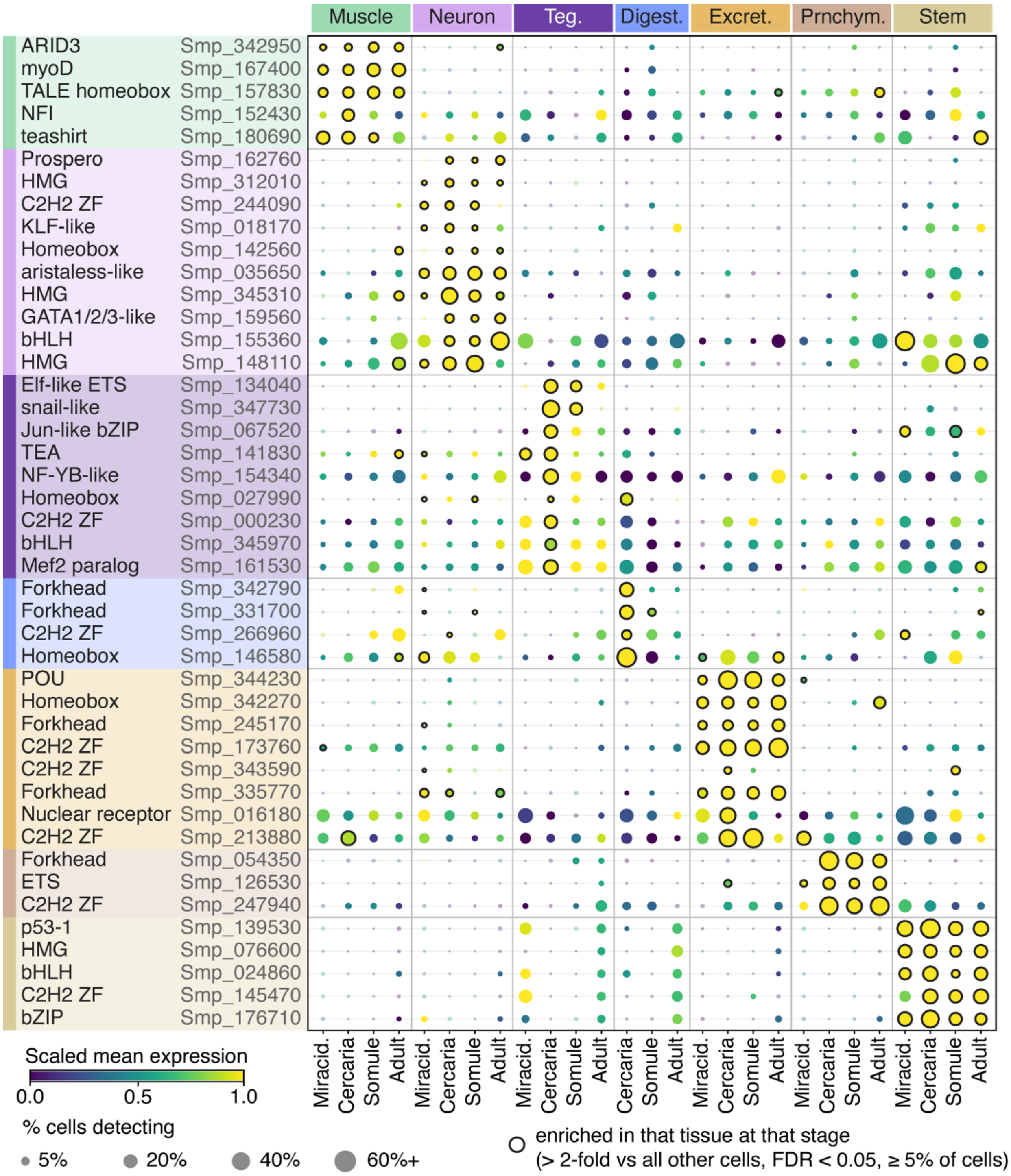


**Fig. S13. Tissue TFs at other life stages.** The tissue TFs of Fig. 4B in the miracidium, cercaria, 2-day schistosomulum and adult atlases, with each atlas's cell types pooled by tissue. Rings, the TF is enriched in that tissue at that stage (cell types of the tissue pooled against all other cells of that atlas, > 2-fold, FDR < 0.05, detected in ≥ 5% of cells of a member cell type), whether its own tissue or another. In its own tissue, 21/36 (miracidium), 31/44 (schistosomulum) and 23/41 (adult) testable TFs were enriched, against ~3 expected by chance. Tests: edgeR, sex pools as replicates for the schistosomulum, libraries for the miracidium and adult. Dot size, % cells detecting; color, mean expression scaled per TF within each stage.

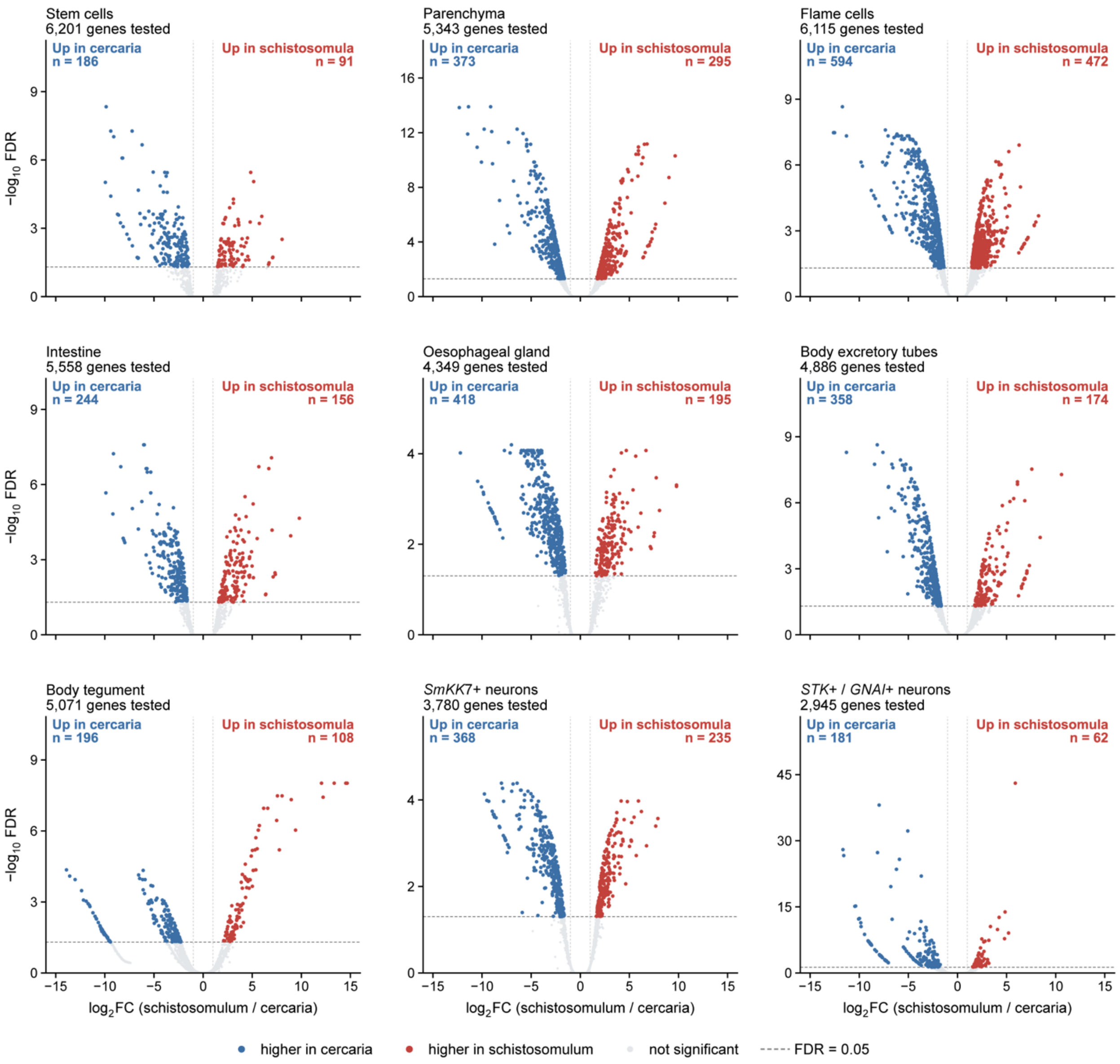


**Fig. S14. Differential expression between cercaria and schistosomulum in each matched cell type.** Volcano plots for the nine cell-type pairs matched between the cercarial and 2-day schistosomulum atlases (table S4). Each point is a gene; the number of genes tested is given above each panel. Blue, higher in the cercaria; red, higher in the schistosomulum; gray, not significant. Tests: edgeR quasi-likelihood against a twofold threshold (glmTreat), with counts summed per sex pool and the two pools as replicates, FDR < 0.05. Dashed line, FDR = 0.05; dotted lines, twofold change. Nine pairs pooled are in Fig. 5A; numbers are in table S11 and gene-level results in data S5.

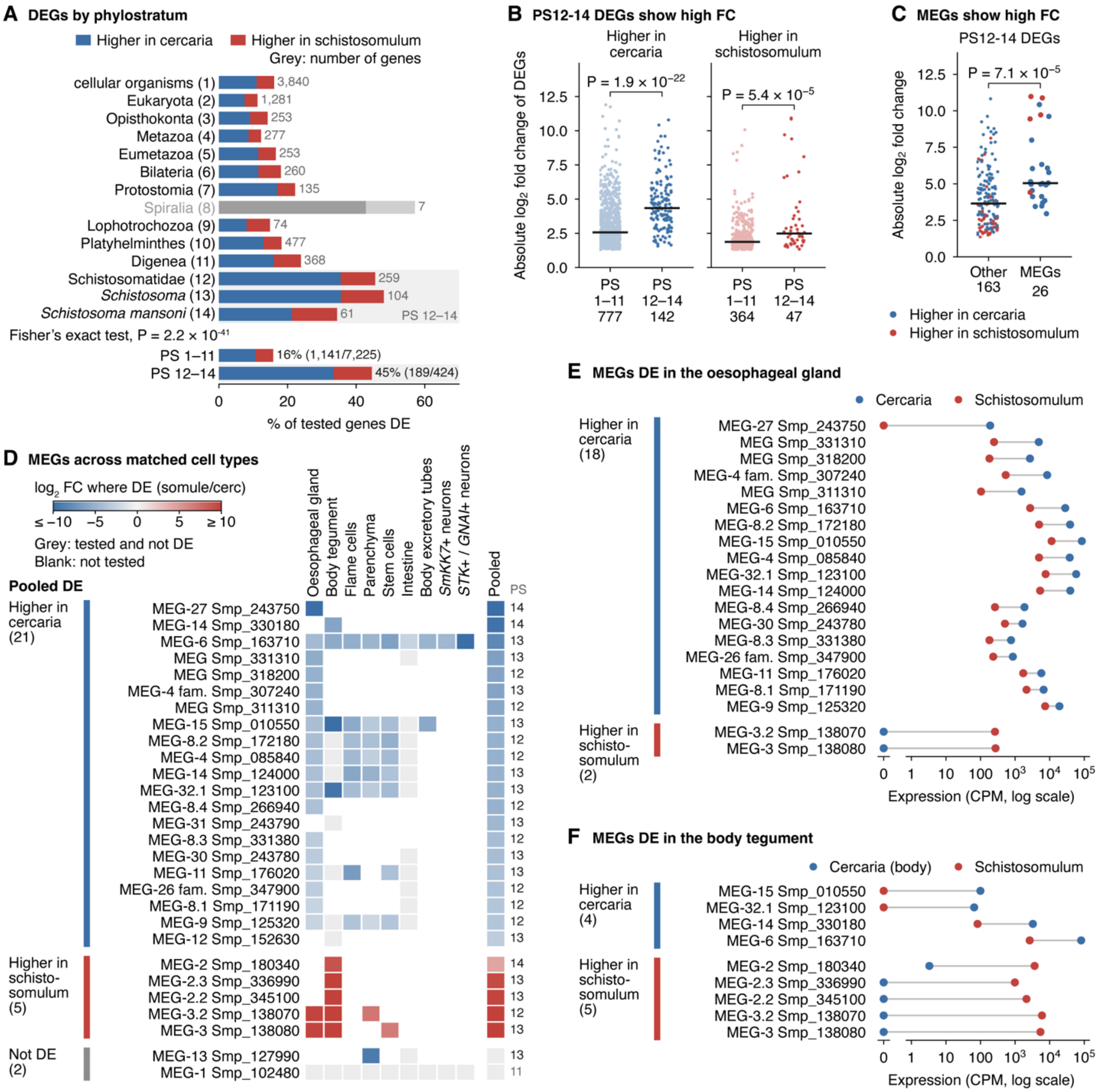


**Fig. S15. Young genes and micro-exon genes (MEGs) across the cercaria–schistosomulum transition. (A)** Percentage of tested genes differentially expressed (DE) in each phylostratum (PS; table S12), pooled comparison of the nine matched cell-type pairs; PS 12–14 (Schistosomatidae and younger) shaded. Spiralia grayed because only seven genes were tested. Bottom, PS 1–11 vs. PS 12–14 (Fisher's exact test). **(B)** Absolute $\log_2$ fold change (FC) of DE genes, PS 1–11 vs. PS 12–14, by direction. **(C)** Absolute $\log_2$ FC of PS 12–14 DE genes, MEGs (table S13) vs. other genes. In (B) and (C), bars show medians; *P* values are from two-sided Mann–Whitney tests. **(D)** The 28 MEGs tested in the pooled comparison, grouped by pooled direction, in each matched pair and pooled. **(E and F)** MEGs DE in the oesophageal gland (E) and body tegument (F): expression in each stage's cell type (counts per million, UMIs summed over libraries; log scale, undetected genes at 0). DE: edgeR, sex pools as replicates, > 2-fold, FDR < 0.05.

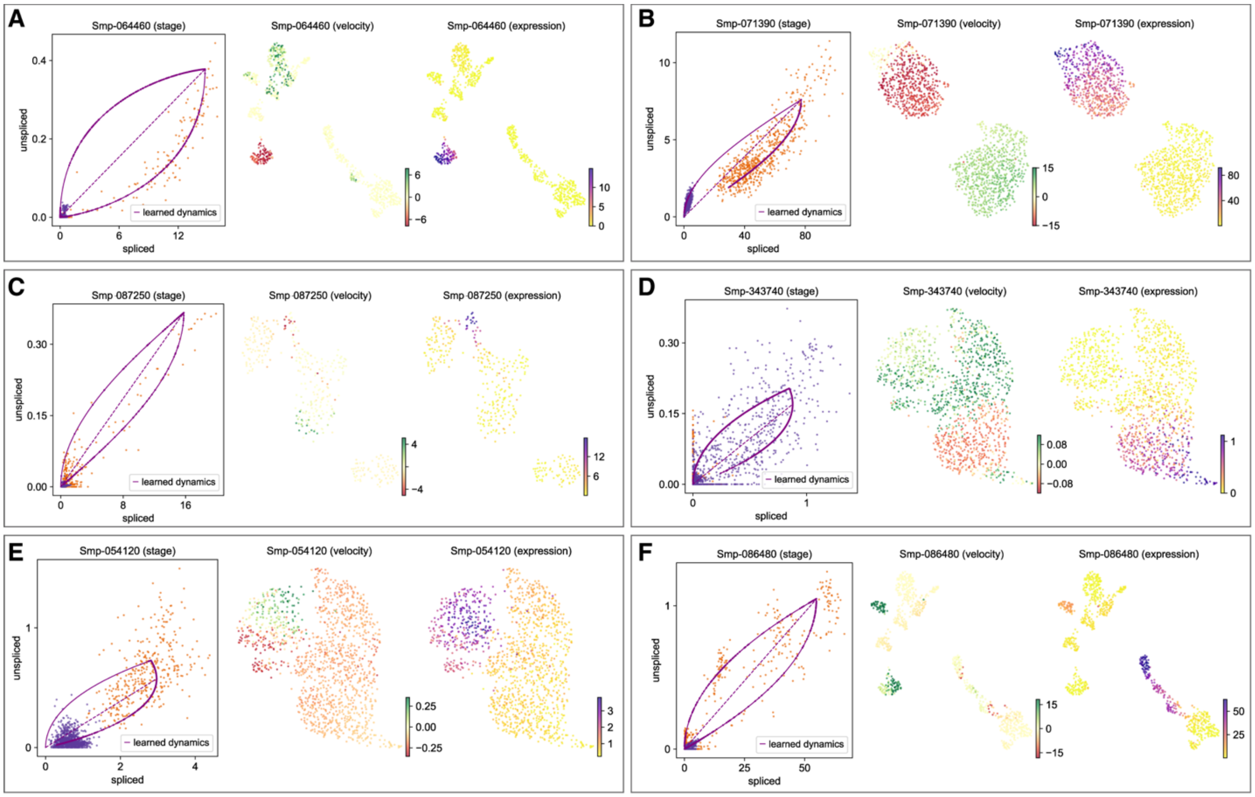


**Fig. S16. RNA velocity.** The transcriptional dynamics between cercariae and schistosomula were analyzed with RNA velocity to quantify the changes and identify important genes in the transition, using the nine cell-type pairs with analogous clusters in both stages. Some genes fit the RNA velocity model well, such as **(A)** *Smp_064460* in the excretory tubes and **(B)** *Smp_071390* in the flame cells. Many genes were a poor fit to the RNA velocity model, such as **(C)** *Smp_078250* in *SmKK7*+ neurons and **(D)** *Smp_343740* in stem cells. Some genes did not fit the model well, and their expression patterns suggest that a different model may apply, such as **(E)** *Smp_054120* in stem cells and **(F)** *Smp_086480* in the excretory tubes. Cercarial cells, orange; schistosomulum cells, purple.

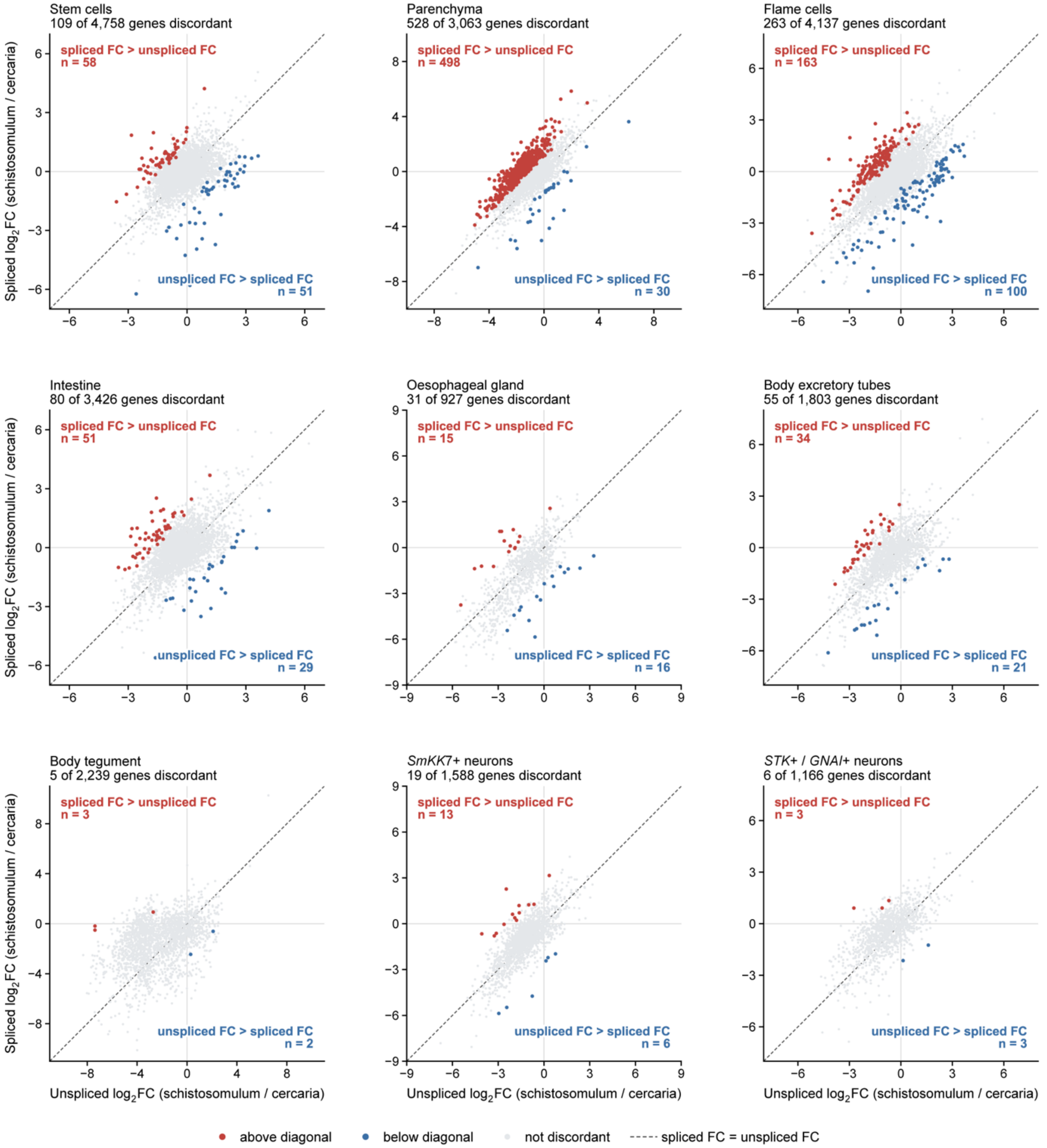


**Fig. S17. Exon–intron split analysis (EISA) in each matched cell type.** Change from cercaria to schistosomulum in unspliced (intronic) reads ($x$) and spliced (exonic) reads ($y$) for each gene, in the nine matched cell-type pairs. Dashed line, equal spliced and unspliced change. Colored points are discordant genes (eisaR, FDR < 0.05). Red, above the diagonal: spliced reads increase more, or decrease less, than unspliced reads in the schistosomulum. Blue, below the diagonal: the reverse. Gray, not discordant. Nine pairs pooled are in Fig. 5D. Gene-level results are in data S6.

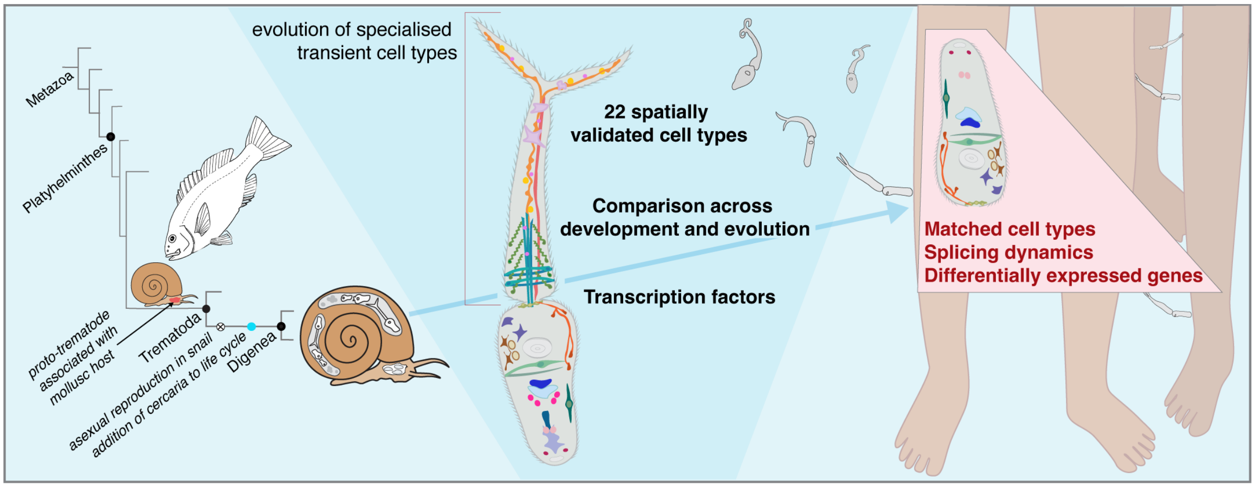


**Fig. S18 Summary of cercaria and schistosomulum atlases in evolutionary context.** In the ancestral digenean, asexual reproduction inside the snail evolved leading to amplification of parasite numbers and the development of the cercaria larva. The shift from ingestion-based transmission of one parasite to one vertebrate host (e.g. a fish) in the proto-trematode to tail-driven active dispersal of many cercariae in the ancestral digenean enabled the chance to infect more than one definitive host. Our atlases show that the cercaria has cellular novelty: its transient tail comprises cell types found nowhere else in the life cycle or in other flatworms, and is built by layering a tail-wide regulatory signature onto existing tissue programmes and drawing disproportionately on young genes. We show that many body cell types persist over the transformation to the schistosomulum, modify their energy metabolism, undergo cell-type-specific reprogramming and have complex transcriptional dynamics.
